# Species-dependent antifungal profiles reveal stronger yeast inhibition by chitosan than by a sulfate-containing polysaccharide-rich extract from *Jania adhaerens*

**DOI:** 10.64898/2026.07.21.739751

**Authors:** Miguel Valverde-Urrea, Jose Defez-Pérez, F. Colom-Valiente, Marc Terradas-Fernández, Luis V. López-Llorca, Federico Lopez-Moya

## Abstract

Yeast infections are becoming an increasing public health concern, mainly due to the spread of opportunistic species and the emergence of strains resistant to commonly used antifungal drugs. Marine resources are a promising source of bioactive compounds, including polysaccharides and other biopolymers with potential antifungal applications. In this study, a sulfate-containing polysaccharide-rich extract was obtained from the red alga *Jania adhaerens* and chemically characterized. Its antifungal activity was compared with that of a commercial chitosan formulation against clinically relevant yeasts, including species of *Candida, Cryptococcus, 'Clavispora, Naganishia* and *Trichosporon*. Growth kinetics were monitored in liquid medium over 24 h, and antifungal activity was evaluated through growth rate analysis, growth inhibition at 20 h and susceptibility clustering. The polysaccharide extract showed moderate but consistent growth inhibition, with the strongest effects observed at 5 mg mL^−1^. Maximum growth inhibition reached 60.9% in *Cryptococcus deuterogattii* and 59.8% in *Candida albicans*, although no complete inhibition was observed within the tested concentration range. In contrast, chitosan showed a stronger antifungal effect, with minimal inhibitory concentration (MIC) values between 10 and 20 µg mL^−1^ in several species and maximum inhibition values above 80% in the most susceptible yeasts. However, *C. albicans* showed marked resistance to chitosan, with inhibition below 12%. K-means clustering confirmed distinct susceptibility profiles between treatments, supporting a species-dependent response. Overall, these results highlight marine-derived biopolymers as promising antifungal candidates and show that chitosan and algal sulphated polysaccharides produce distinct, species-dependent antifungal profiles.

## 1. Introduction

Fungal infections are an increasing global health problem, with more than 1.5 million deaths reported annually [1,2]. Although more than 1.5 million fungal species are estimated to exist, only ca. 600 can infect humans [3,4]. Among these, *Aspergillus, Candida* and *Cryptococcus* spp. are the most clinically relevant [5]. Furthermore, *Candida* spp. Are responsible for the majority of human mycoses [6–8].

Infections caused by *Candida* species are of particular concern due to their high incidence and associated healthcare costs. These infections range from superficial or mucosal candidiasis to invasive disease, including candidemia, which refers to the presence of *Candida* in the bloodstream. The estimated incidence of candidiasis ranges from 2 to 14 cases per 100,000 inhabitants, while candidemia affects up to 250,000 people annually [9]. Among these species, *Candida auris* has emerged as a major pathogenic yeast threat due to its resistance to multiple antifungal drug classes, including polyenes, azoles and echinocandins [10,11]. *C*.*auris* has been reported in more than 30 countries and is associated with high mortality rates and environmental persistence, which facilitates horizontal transmission [8,12,13]. Beyond *Candida*, other opportunistic yeasts, particularly species of *Cryptococcus, Clavispora, Naganishia* and *Trichosporon*, are also increasingly recognized as clinically relevant pathogens.

Infections caused by *Cryptococcus* species (cryptococcosis) mainly affect immunocompromised patients, including those with HIV infection, diabetes or lymphocytopenia, as well as individuals undergoing treatment with corticosteroids or immunosuppressive drugs [14–15]. Other emerging or opportunistic yeasts also represent a growing concern, such as *Clavispora lusitaniae*. This yeast can cause severe infections such as meningitis and peritonitis and shows resistance to commonly used antifungal agents such as fluconazole and amphotericin B [16]. Although *Naganishia albida* is not typically pathogenic, increasing cases of infection in immunocompromised patients have been reported [17,18]. Similarly, *Trichosporon cutaneum* is considered an opportunistic emerging pathogen, often associated with biofilm formation on medical devices, which facilitates cross-infections [19].

Despite the availability of antifungal drugs, the emergence of resistant strains is increasing and represents a major clinical challenge [20]. This situation highlights the need to identify new antifungal compounds and alternative therapeutic strategies [21].

Marine resources constitute a promising source of bioactive molecules due to their high diversity, abundance and unique chemical structures [22–25]. Among these resources, marine algae have attracted considerable attention. Several studies have demonstrated the antifungal activity of algal extracts against human pathogenic fungi. For example, [26] reported growth inhibition of *Candida* species using polar extracts from *Saccharina japonica, Laminaria japonica, Undaria pinnatifida, Palmaria palmata* and *Eisenia bicyclis*. In addition, apolar extracts from *Jania adhaerens* have shown antifungal activity against *C. albicans* [27].

Among algal-derived compounds, sulphated polysaccharides (PS) have gained particular interest due to their wide range of biological activities, including antifungal properties [28,29]. These compounds are characterized by the presence of sulphate groups (SO_3_^−^) attached to sugar residues. Algae represent one of the main natural sources of PS, including fucoidans, carrageenans, galactans and agar. The composition of these polysaccharides depends on the algal group, with sulphated galactans predominating in red algae [30]. Although some studies have reported antifungal effects of PS, such as the alteration of fungal cell morphology by carrageenans in *Aspergillus fumigatus* [31], their activity against clinically relevant yeasts such as *Candida* and *Cryptococcus* remains poorly characterized [32].

Articulated coralline algae are dominant along the Alicante coast and show high tolerance to environmental stress [33,34]. Among them, *Jania adhaerens* is abundant and easily collected, forming conspicuous belts along the coastline [35]. This species has shown high polysaccharide extraction yields and relevant sulfate content [30], making it a suitable candidate for antifungal studies.

In addition to algal-derived compounds, other marine biopolymers such as chitosan have shown strong antifungal activity [36,37]. Chitosan, the N-deacetylated form of chitin, exhibits favourable physicochemical properties, including solubility in mildly acidic conditions and a positive charge that enhances its interaction with microbial cells [38]. It is mainly obtained from seafood biomass, contributing to circular economy strategies and reducing environmental impact [39,40].

The antifungal activity of chitosan has been widely studied. [41] reported fungistatic effects on *C. albicans* at low concentrations (2.5–10 µg mL^−1^), likely related to membrane disruption, ion imbalance and metabolic alterations. Other mechanisms, such as metal chelation, enzyme inhibition and interaction with DNA, have also been proposed [42]. In addition, chitosan has shown activity against other clinically relevant yeasts, including *C. glabrata, C. parapsilosis* and *Cryptococcus* species [43], even in fluconazole-resistant strains [44].

However, comparative studies evaluating different marine-derived biopolymers across a wide range of clinically relevant yeast species remain scarce. In addition, the antifungal activity of algal sulphated polysaccharides have been poorly explored in emerging pathogenic genera such as *Clavispora, Naganishia* and *Trichosporon*.

In this context, the present study aims to evaluate the antifungal activity of marine derivates mainly focused on polysaccharides extracted from *Jania adhaerens* and crustacean derived chitosan against important human pathogenic yeasts, contributing to the identification of marine-derived bioactive compounds for antifungal applications.

## 2. Materials and Methods

### 2.1 Yeast strains

The antifungal activity of algal polysaccharides was tested against the following species: *Candida albicans, C. auris, Cryptococcus bacillisporus* and *Cr. deuterogattii*. For chitosan assays, a broader set of species was used: *Candida albicans, C. auris, C. glabrata, C. guilliermondii, C. parapsilosis, Cryptococcus bacillisporus, Cr. deuterogattii, Cr. deneoformans, Cr. tetragattii, Clavispora lusitaniae, Naganishia albida* and *Trichosporon cutaneum*. All yeast strains were kindly provided by Dr. Colom-Valiente from the Plant Production and Microbiology Laboratory (Miguel Hernández University, Spain).

### 2.2 Chitosan formulation

Liquid chitosan was prepared from commercial chitosan powder (T8s, Biolog Hepe, Germany). The chitosan used had a medium molecular weight (70 kDa) and a degree of deacetylation of 90.1%. The powder was dissolved in 0.25 M hydrochloric acid (HCl), and the pH was adjusted to 5.60 [45]. The solution was dialyzed for 48 h using a cellulose membrane (12–15 kDa cut-off; Sigma-Aldrich, USA) to remove salts. Finally, the solution was sterilized at 121 °C for 20 min. The liquid chitosan was stored at 4 °C until use.

### 2.3 Algae biomass collection and morphological and molecular identification

Algal biomass was collected at Agua Amarga beach (Alicante, Spain; 38.299237°, −0.518894°), kept on ice and protected from light until processing. Samples were cleaned and morphologically identified using a Leica MZ6 stereomicroscope and an Olympus BH-2 microscope equipped with a Leica DFC camera [46].

For molecular identification, DNA was extracted using a CTAB-based buffer (0.1 M Tris-HCl, 0.05 M EDTA, 1.5 M NaCl, 0.05 M DTT and 3% CTAB) [47].

The cytochrome oxidase I (COI) gene was amplified using primers GazF1 and GazR1 [48]. PCR conditions consisted of an initial denaturation at 94 °C for 4 min, followed by 40 cycles (94 °C for 1 min, 50 °C for 30 s, and 72 °C for 1 min), and a final extension at 72 °C for 7 min. PCR products were visualized on 1% agarose gel, purified (GeneJET Extraction Kit, ThermoScientific, USA) and sequenced by Sanger method (STAB Vida, Portugal).

Consensus sequences were generated using BioEdit v7.2.5 [49] and compared with the NCBI database using megaBLAST (accessed February 10, 2024).

### 2.4 Extraction of algal polysaccharides

Lyophilized and ground algal biomass was treated with 80% ethanol at 80 °C for 4 h to remove pigments, phenols, proteins and salts [50,51]. This step was repeated until complete pigment removal. The sample was filtered and dried, then dissolved in ultrapure water (1:30, w/v) and heated at 100 °C for 1 h [50,51]. After filtration and centrifugation (4000 g, 10 min, 4 °C), the supernatant was dialyzed for 48 h using a 10 kDa cut-off membrane. Polysaccharides were precipitated by adding two volumes of 95% ethanol and incubating at −20 °C for 48 h [51]. The solution was centrifuged (4000 g, 10 min, 4 °C), and the pellet was lyophilized and resuspended in sterile water to obtain a final stock of 10 mg mL^−1^. Extraction yield (%) was calculated as (dry weight of polysaccharides / dry weight of algae) × 100.

#### 2.5.1 Characterization of polysaccharides

The extract was analyzed for lipids, soluble and insoluble carbohydrates, proteins, and mineral content (Ca, Mg, K, Na and S), as well as total sulphated polysaccharides and sulphur content. Lipids were extracted using a Soxhlet system with petroleum ether (40–60 °C) following [52]. Carbohydrates were determined using the anthrone method distinguishing soluble and insoluble fractions [53]. Sulphated polysaccharides were quantified using a turbidimetric method [54]. Protein content was determined by Bradford assay [55]. Elemental composition (Ca, Mg, Na and K) was analyzed by ICP-OES (Perkin Elmer 7300DV). FTIR analysis was performed using a BRUKER FT/IR-4700 spectrophotometer (500–4000 cm^−1^, ATR mode).

#### 2.5.2 Monosaccharide composition

Polysaccharides (10 mg) were hydrolyzed with 6 M HCl at 110 °C for 6 h. After solvent evaporation, samples were resuspended in ultrapure water (100 µg mL^−1^) and filtered (0.22 µm). Monosaccharides were analyzed by anion-exchange chromatography with pulsed amperometric detection.

### 2.6 Antifungal activity assays

Antifungal activity was evaluated by growth kinetics in liquid medium. Yeasts were grown on Sabouraud medium for 24 h at 36 ± 1 °C. A single colony was then transferred to YPD medium (yeast extract, peptone, dextrose) and incubated under the same conditions. Cell suspensions were adjusted to OD490 < 0.1 [56]. All treatments were prepared in 50% YPD medium.

Polysaccharides were tested at 1, 2, 3 and 5 mg mL^−1^, and chitosan at 2.5, 5, 10, 15 and 20 µg mL^−1^. Cultures were incubated for 24 h at 36 ± 1 °C, and optical density was measured hourly at 490 nm using an Infinite F Nano multimode microplate reader (Tecan Trading AG, Zurich, Switzerland). All assays were performed in three independent biological experiments, with three technical replicates per treatment in each experiment.

### 2.7 Statistical analysis

Growth curves were analyzed by fitting an exponential model: N(t) = N_0_e⍰ ⍰, where r is the growth-rate parameter. The model was fitted using the Levenberg–Marquardt algorithm (minpack.lm package [57]). Minimum inhibitory concentration (MIC) was determined both graphically and statistically from r values. Differences between treatments were analyzed by one-way ANOVA. Growth inhibition (%) was calculated at 20 h relative to control. Normality and homoscedasticity were tested using Shapiro–Wilk and Levene tests, respectively, and Tukey HSD was used for post hoc comparisons. K-means clustering was used to group species based on sensitivity, and the optimal number of clusters was determined using silhouette analysis. Principal component analysis was used to visualize the K-means clusters in two-dimensional space. All analyses were performed in R v4.3.3 [58].

A schematic representation of the statistical workflow used in this study is provided in Figure S1.

## 3. Results

### 3.1 Algae identification and characterization

The collected algal biomass was morphologically identified as *Jania adhaerens* based on its characteristic dichotomous thallus, pale pink-lilac coloration and articulated calcified structure (Figure S2). The specimens presented an approximate length of 5 cm, with calcified intergenicula measuring 80–120 µm in diameter and lengths approximately 5–6 times greater than their width. In addition, dichotomy angles were consistently greater than 30°, in agreement with the diagnostic morphological features previously described for *J. adhaerens* (Figure S2). Molecular identification based on COI sequencing further supported this classification, showing ≥99% similarity with reference sequences available in the NCBI database.

### 3.2 *Jania adhaerens* yielded a sulfate-containing polysaccharide extract with characteristic FTIR signatures

Polysaccharide extraction yielded 3.7 ± 0.3% (w/w) relative to the initial dry biomass. The extract was predominantly composed of carbohydrate fractions, with insoluble polysaccharides accounting for 56.23% and soluble polysaccharides for 20.24% of the total extract. Lipids (10.67%) and proteins (2.25%) were present in lower proportions.

Elemental analysis indicated the presence of mineral components, including Ca (2.5%), Mg (0.41%), K (0.028%), Na (0.078%) and S (1.87%).

Monosaccharide analysis revealed a heterogeneous composition dominated by arabinose (62.1%), followed by xylose (17.1%) and fructose (16.0%), while rhamnose was detected at lower proportions (4.7%). This heterogeneous monosaccharide profile is consistent with the structural complexity commonly described for sulphated polysaccharides from red macroalgae. Galactose was not quantified because an analytical standard was not available.

FTIR analysis revealed characteristic absorption bands associated with sulphated polysaccharides (Figure 1). Bands detected between 500–600 cm^−1^ and at 931 cm^−1^ were associated with sulfate groups (S=O and SO_3_^−^), while signals between 1220–1270 cm^−1^ corresponded to sulfate ester bonds (O–SO_3_^−^), which are typical features of sulphated polysaccharides from red algae [59,60]. In addition, the band at 1644 cm^−1^ was related to C=O vibrations, whereas signals observed between 2833–2963 cm^−1^ and 3412–3433 cm^−1^ were attributed to C–H and O–H stretching vibrations, respectively, confirming the polysaccharidic nature of the extract [60,61]. Sulfate groups were detected using a turbidimetric method, estimating a sulfate content of 11.3% of the dry extract.

**Figure 1.**
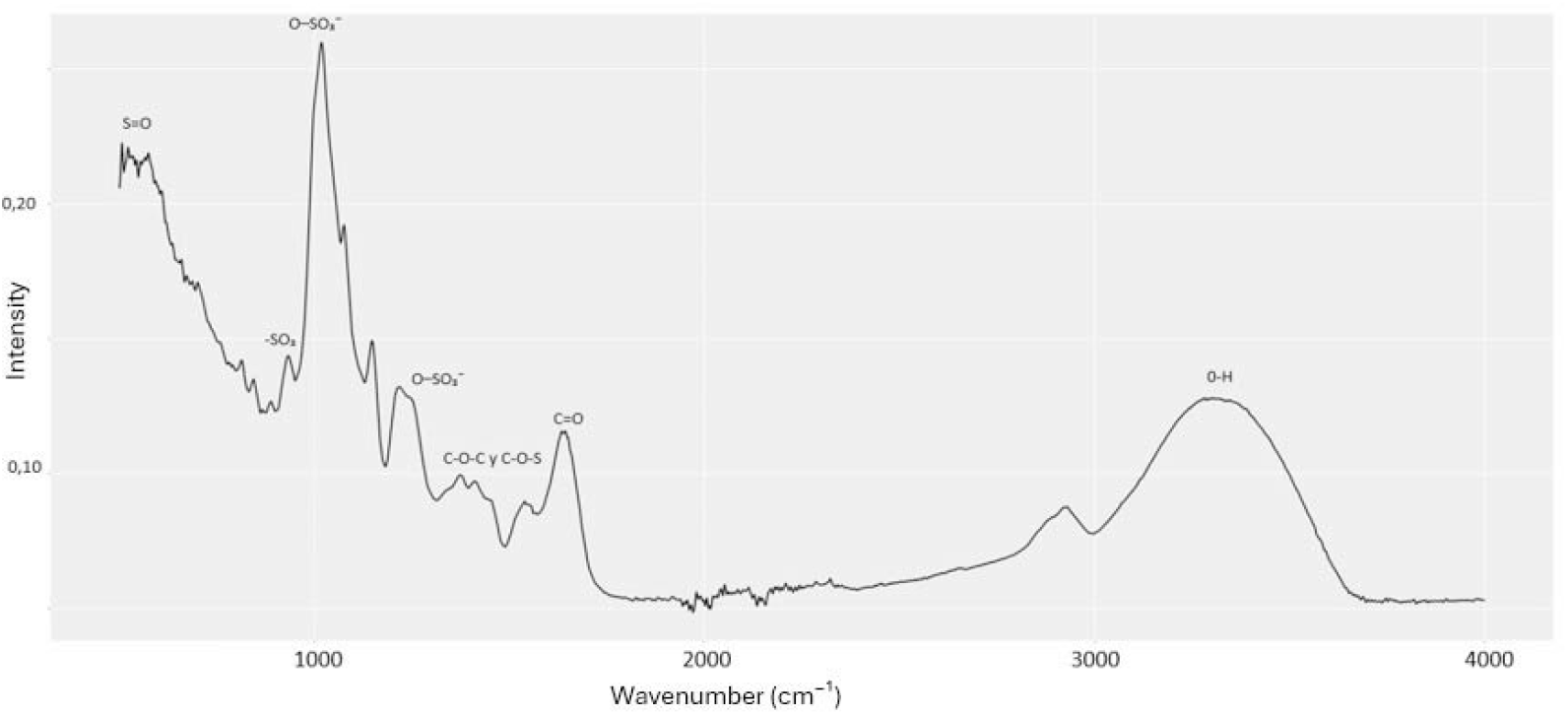
FTIR analysis of the crude polysaccharide extract obtained from *J. adhaerens*.

### 3.3 Chitosan showed strong but species-dependent antifungal activity against pathogenic yeasts

Chitosan showed a marked antifungal effect against most of the yeast species tested, although the response strongly depended on both species and concentration (Figures S3-S5). Growth kinetics and growth rate parameter (r) analyses revealed a progressive reduction in fungal growth as chitosan concentration increased, with significant differences detected in all species (ANOVA, p < 0.05; Figures S3–S11, Tables S1-S3). However, susceptibility patterns varied considerably among species, ranging from highly sensitive to moderately resistant responses.

The highest antifungal activity was observed in *Cr. deneoformans, Cl. lusitaniae, Cr. tetragattii, C. parapsilosis* and *N. albida*, which reached maximum inhibition values above 80% and MIC values between 10 and 20 µg mL^−1^ (Table 1, Figure 2). In contrast, *C. albicans* showed very low sensitivity to chitosan, with inhibition values remaining below 12% even at the highest concentration tested, despite a significant reduction in the growth rate parameter. Similarly, *T. cutaneum* and *C. guilliermondii* maintained detectable growth at all tested concentrations, indicating lower susceptibility to the treatment.

**Table 1.** Minimum inhibitory concentration (MIC) and maximum growth inhibition values of chitosan and the polysaccharide-rich extract (PS) extracted from *Jania adhaerens* against different yeast species. MIC values are expressed as µg mL^−1^ for chitosan and mg mL^−1^ for polysaccharides-rich extract. Maximum growth inhibition refers to the highest percentage reduction in OD_490_ at 20 h relative to the untreated control. Values are presented as mean ± standard deviation. † No detectable increase in OD_490_ was observed during the assay, indicating apparent growth suppression under the tested conditions.

| Species | MIC chitosan ($\mu\text{g mL}^{-1}$) | MIC PS (mg $\text{mL}^{-1}$) | Max inhibition chitosan (%) | Max inhibition PS (%) |
| --- | --- | --- | --- | --- |
| *C. albicans* | >20 | >5 | 11.5 ± 3.21 | 59.8 ± 2.69 |
| *C. auris* | 20 | >5 | 78.2 ± 1.26† | 55.02 ± 7.01 |
| *C. glabrata* | 20 | - | 69.8 ± 0.74† | - |
| *C. guilliermondii* | >20 | - | 72.2 ± 0.28 | - |
| *C. parapsilosis* | 15 | - | 82.6 ± 0.32† | - |
| *Cr. bacillisporus* | 10 | >5 | 76.7 ± 0.87† | 30.39 ± 3.55 |
| *Cr. deneoformans* | 15 | - | 87.7 ± 1.04† | - |
| *Cr. deuterogattii* | 15 | >5 | 74.8 ± 3.09† | 60.88 ± 1.94 |
| *Cr. tetragattii* | 20 | - | 85.1 ± 0.35† | - |
| *Cl. lusitaniae* | 20 | - | 87.0 ± 0.26† | - |
| *N. albida* | 20 | - | 81.2 ± 0.55† | - |
| *T. cutaneum* | >20 | - | 69.4 ± 3.59 | - |

**Figure 2.**
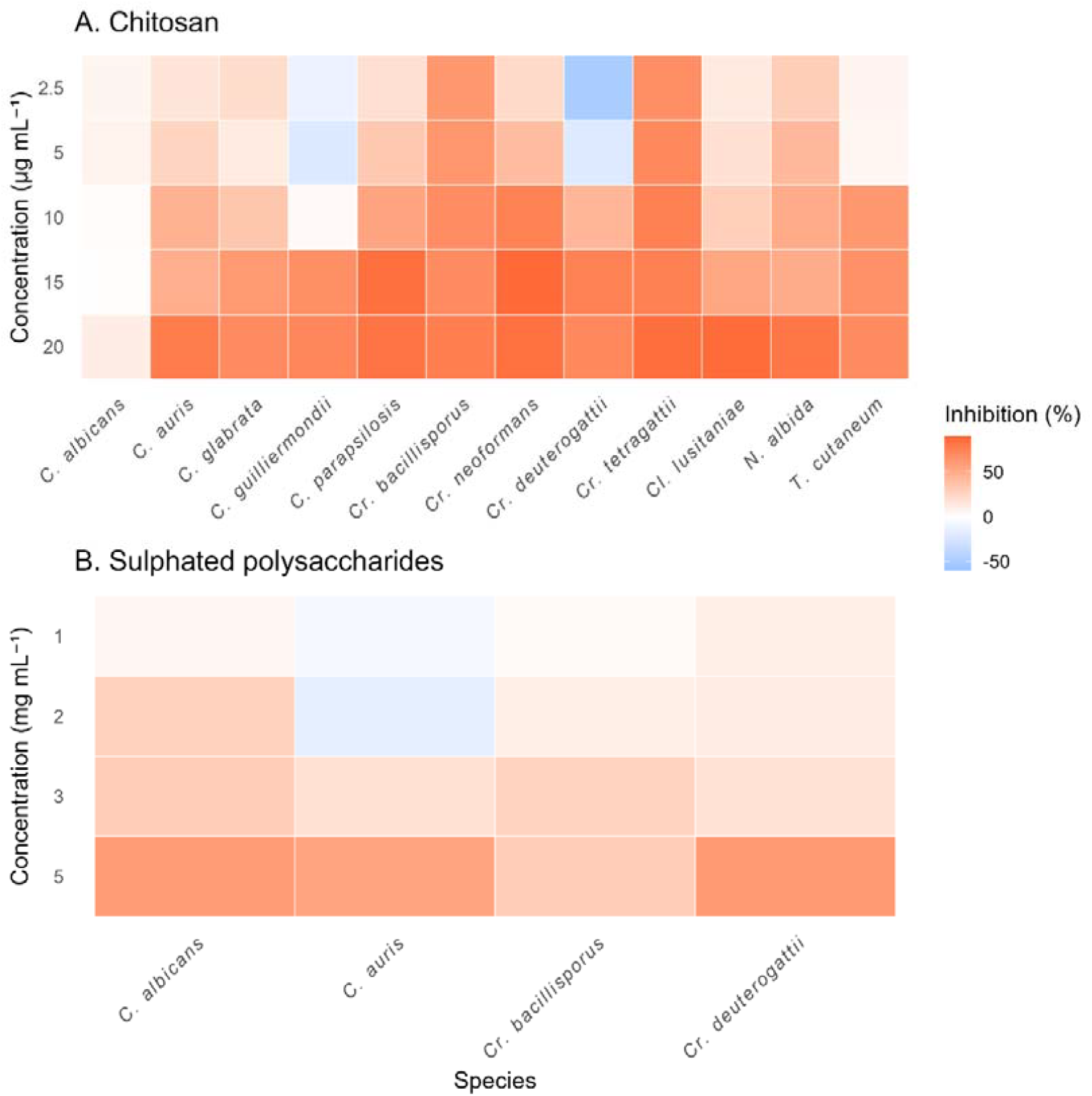
Heatmap representation of the antifungal activity of (A) chitosan and (B) polysaccharide-rich extract from *J. adhaerens* against different yeast species at increasing concentrations. Colours represent the percentage of growth inhibition relative to the untreated control, ranging from stimulation (blue tones, negative values) to strong inhibition (orange-red tones, positive values).

Interestingly, some species, particularly *C. guilliermondii* and *Cr. deuterogattii*, showed growth stimulation at low chitosan concentrations before inhibition occurred at higher doses. Overall, growth inhibition generally increased in a concentration-dependent manner, reaching values between 69% and 88% in the most susceptible species.

### 3.4. Polysaccharides-rich extract showed weaker but species-dependent growth inhibition compared with chitosan

The polysaccharide-rich extract showed a moderate and species-dependent antifungal effect (Figures S12–S14). Growth rates differed significantly among treatments (one-way ANOVA, p < 0.05, Tables S4-S5), although the magnitude of the effect was lower compared to chitosan.

No complete inhibition was observed within the tested concentration range (1–5 mg mL^−1^), and MIC values were consistently above 5 mg mL^−1^ (Table 1, Figure 2).

Maximum inhibition values reached 60.9% in *Cr. deuterogattii* and 59.8% in *C. albicans* at 5 mg mL^−1^, whereas other species, such as *Cr. bacillisporus*, showed lower inhibition levels, not exceeding 30%.

As observed for chitosan, low concentrations occasionally resulted in slight growth stimulation. This effect was particularly evident in *C. auris*, where negative inhibition values were recorded at 1 and 2 mg mL^−1^.

### 3.5 K-means clustering revealed distinct susceptibility profiles between treatments

To further explore species-specific responses to both treatments, a K-means clustering analysis was performed based on inhibition profiles across concentrations (Figure 3).

**Figure 3.**
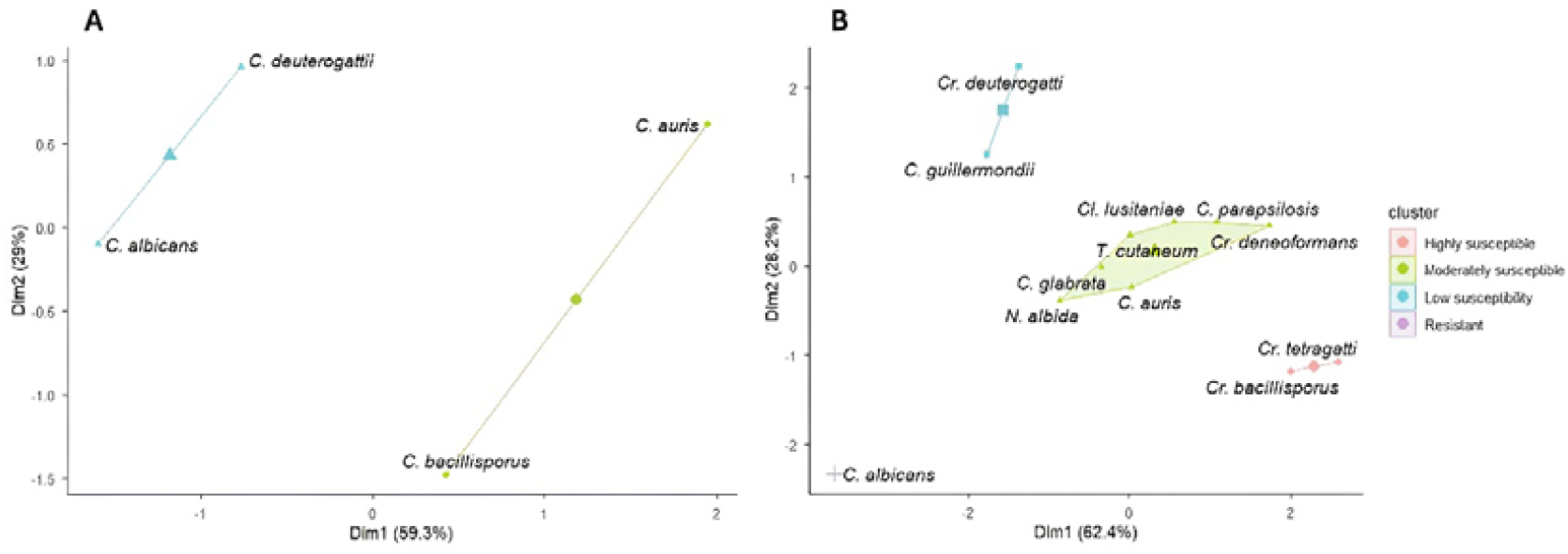
K-means clustering analysis of the antifungal response patterns of different yeast species treated with (A) the polysaccharide-rich extract obtained from *Jania adhaerens* and (B) commercial chitosan. Species were grouped according to their inhibition profiles across the tested concentration ranges.

For chitosan-treated yeast, the analysis revealed a clear separation of species according to their sensitivity to chitosan. Highly sensitive species, mainly within the genus *Cryptococcus*, showed strong inhibition even at low concentrations, whereas other species displayed a more gradual, dose-dependent response. In contrast, *Candida albicans* showed minimal inhibition across all concentrations and was clearly separated from the rest of the dataset. Intermediate patterns were observed in species such as *C. auris* and *C. parapsilosis*, which exhibited a progressive increase in inhibition with increasing chitosan concentration.

Some species did not follow a simple dose–response pattern. In particular, *Cr. deuterogattii* and *C. guilliermondii* showed a biphasic behaviour, with stimulation of growth at low concentrations followed by inhibition at higher doses. This pattern contributed strongly to their position in the multivariate space and highlights the variability in the response to chitosan among yeast species.

For polysaccharide-rich extract, clustering showed a more limited differentiation between species, with most responses grouped within a narrower range of inhibition values. Although inhibition increased with concentration, the overall effect was lower compared to chitosan, and no species reached complete inhibition within the tested range.

The first principal component explained more than 60% of the total variance, indicating that the main trends in the data were well captured by the analysis. Overall, clustering supported the existence of distinct response patterns between species and confirmed the stronger and more variable antifungal effect of chitosan compared to polysaccharide-rich extract.

## 4. Discussion

The polysaccharide extraction yield obtained from *J. adhaerens* (3.7 %) was consistent with the value previously reported by Hentati et al. [30] (4.55%). This close agreement was observed despite the additional washing steps applied in the present study to remove pigments, phenolic compounds, proteins, and salts, which were intended to improve the purity and suitability of the extract for subsequent chemical and biological characterization.

Despite the lower yield, the extracted polysaccharides showed a consistent inhibitory effect on all tested yeast species at 5 mg mL^−1^. This is the first study reporting antifungal activity of polysaccharides from *J. adhaerens* against clinically relevant yeasts. However, the observed effect was moderate compared to chitosan, suggesting a lower intrinsic antifungal potency or a different mode of action. The presence of sulphate groups and sulphur content indicates that these compounds are likely sulphated polysaccharides, potentially belonging to the group of sulphated galactans typically found in red algae, although further structural characterization would be required to confirm their precise nature.

The mechanism underlying the antifungal activity of these polysaccharides remains unclear. Previous studies have reported alterations in fungal cell wall composition following exposure to algal polysaccharides, including reductions in β-1,3-glucan and chitin content [31]. In addition, sulphated polysaccharides have been associated with DNA damage, receptor interactions and metabolic disruption in microbial systems [63,64]. Their ability to interfere with biofilm formation and gene expression has also been described [65]. Although these mechanisms have mainly been reported in bacteria or filamentous fungi, they could also contribute to the inhibitory effects observed in yeasts.

In contrast, chitosan exhibited strong antifungal activity across most species, with low MIC values (10–20 µg mL^−1^), confirming its broad antifungal activity across the tested species. This result is consistent with previous studies describing chitosan as a potent antifungal agent in both yeast and filamentous fungi [41, 43]. The antifungal activity of chitosan has been attributed to its cationic nature, which enables interaction with negatively charged membrane phospholipids, leading to membrane disruption and increased permeability [45,66,67]. In addition, transcriptional changes affecting cell wall synthesis and stress response pathways have been reported [68, 69, 72].

Differences in antifungal activity between studies may be explained by experimental conditions. In particular, nutrient availability has been shown to modulate sensitivity to chitosan, with carbon and nitrogen limitation increasing antifungal susceptibility [43]. The relatively high nutrient content of the medium used in this study may therefore explain the reduced sensitivity observed in *Candida albicans*, which was the most resistant species.

The physicochemical properties of chitosan, including molecular weight and degree of deacetylation, also play a key role in its antifungal activity. In this study, chitosan with a relatively high molecular weight (70 kDa) and a high degree of deacetylation (90.1%) was used, both of which have been associated with increased biological activity [70,71]. Moreover, fungal sensitivity to chitosan depends on membrane composition and cell wall structure, particularly the glucan-to-chitin ratio and fatty acid profile [45,72]. The higher resistance of *C. albicans* observed in this study may therefore be related to its membrane composition, which is enriched in saturated fatty acids [73].

Interestingly, *Candida auris* showed sensitivity to chitosan at 20 µg mL^−1^, highlighting its potential relevance as an alternative antifungal agent against this multidrug-resistant pathogen [74,75]. Given the limited therapeutic options currently available for *C. auris*, these results support further exploration of chitosan-based strategies. Although direct comparisons with conventional antifungals were not performed in this study, the observed response supports further evaluation of chitosan-based strategies against this multidrug-resistant pathogen. Moreover, chitosan can be obtained from marine by-products such as crustacean waste, making it an environmentally friendly biopolymer aligned with circular economy approaches.

Another relevant observation was the stimulation of growth at low concentrations of both chitosan and polysaccharides in some species. This effect may be related to hormetic responses or to the antioxidant properties of chitosan [76], which could enhance cell survival under mild stress conditions.

A key finding of this study is the differential response of yeast species to chitosan and sulphated polysaccharides. For instance, *C. albicans* showed resistance to chitosan but remained moderately sensitive to polysaccharides. This contrast likely reflects differences in their mechanisms of action. Chitosan, as a cationic polymer, primarily targets membrane integrity, whereas sulphated polysaccharides, being anionic, may interact differently with the cell surface and affect biofilm formation or cellular metabolism [65,68]. These complementary modes of action suggest that combined treatments could enhance antifungal efficacy, an approach that warrants further investigation.

Future studies should explore the influence of nutrient availability on antifungal activity, as well as evaluate these compounds under conditions more closely resembling physiological environments. In addition, testing drug-resistant strains and assessing potential synergistic effects between T8 chitosan and algal sulfated polysaccharides could provide valuable insights into their applicability as alternative or complementary antifungal strategies.

## 5. Conclusions

Chitosan and polysaccharides extracted from *Jania adhaerens* exhibited antifungal activity against a broad range of clinically relevant yeasts. However, antifungal efficacy strongly depended on both antifungal concentration and target species, highlighting marked interspecific variability in susceptibility. Chitosan showed higher potency at low concentrations, whereas algal polysaccharides displayed a moderate but consistent inhibitory effect.

The contrasting responses observed between treatments suggest distinct mechanisms of action, which may offer complementary antifungal strategies. In particular, the sensitivity of multidrug-resistant species such as *Candida auris* to chitosan supports further evaluation of chitosan against *Candida auris*..

Overall, these findings support marine-derived biopolymers as promising candidates for antifungal applications and encourage further studies to evaluate their mechanisms and potential combined use.

## Author contributions (CRediT)

Valverde-Urrea, M. contributed to conceptualization, methodology, investigation, formal analysis, and writing – original draft. Défez-Pérez, J. contributed to investigation and data curation. Colom-Valiente, F. contributed to conceptualization, supervision, and writing – review & editing. Terradas-Fernández, M. contributed to writing – review & editing. Lopez-Llorca, L. V. contributed to supervision and writing – review & editing. Lopez-Moya, F. contributed to conceptualization, supervision, and writing – review & editing. All authors contributed to data interpretation and approved the final version of the manuscript.

## Data availability

The datasets generated and/or analyzed during the current study are available from the corresponding author upon reasonable request.

## References

[1] Bongomin, F.; Gago, S.; Oladele, R.O.; Denning, D.W. Global and multi-national prevalence of fungal diseases—estimate precision. J. Fungi 2017, 3, 57. 10.3390/jof3040057

[2] World Health Organization. WHO Fungal Priority Pathogens List to Guide Research, Development and Public Health Action; World Health Organization: Geneva, Switzerland, 2022.

[3] Hawksworth, D.L.; Lücking, R. Fungal diversity revisited: 2.2 to 3.8 million species. Microbiol. Spectr. 2017, 5, 1–17. 10.1128/microbiolspec.FUNK-0052-2016

[4] Fisher, M.C.; Henk, D.A.; Briggs, C.J.; Brownstein, J.S.; Madoff, L.C.; McCraw, S.L.; Gurr, S.J. Emerging fungal threats to animal, plant and ecosystem health. Nature 2012, 484, 186–194. 10.1038/nature10947

[5] Bassetti, M.; Peghin, M.; Timsit, J.F. The current treatment landscape: candidiasis. J. Antimicrob. Chemother. 2016, 71, ii13–ii22. 10.1093/jac/dkw392

[6] Singh, S.; Vijayaraghavan, P.; Devi, S.; Hameed, S. An update on human fungal diseases: A holistic overview. In Advances in Antifungal Drug Development: Natural Products with Antifungal Potential; Springer: Singapore, 2024; pp. 3–37. 10.1007/978-981-97-5165-5_1

[7] Pappas, P.G.; Lionakis, M.S.; Arendrup, M.C.; Ostrosky-Zeichner, L.; Kullberg, B.J. Invasive candidiasis. Nat. Rev. Dis. Primers 2018, 4, 18026. 10.1038/nrdp.2018.26

[8] Bays, D.J.; Jenkins, E.N.; Lyman, M.; Chiller, T.; Strong, N.; Ostrosky-Zeichner, L.; Hoenigl, M.; Pappas, P.G.; Thompson, G.R., III. Epidemiology of invasive candidiasis. Clin. Epidemiol. 2024, 16, 549–566. 10.2147/CLEP.S459600

[9] Kullberg, B.J.; Arendrup, M.C. Invasive candidiasis. N. Engl. J. Med. 2015, 373, 1445–1456. 10.1056/NEJMra1315399

[10] Rhodes, J.; Fisher, M.C. Global epidemiology of emerging Candida auris. Annu. Rev. Microbiol. 2019, 73, 429–449. 10.1146/annurev-micro-020518-115635

[11] Ademe, M.; Girma, F. Candida auris: From multidrug resistance to pan-resistant strains. Infect. Drug Resist. 2020, 13, 1287–1294. 10.2147/IDR.S249864

[12] Forsberg, K.; Woodworth, K.; Walters, M.; Berkow, E.L.; Jackson, B.; Chiller, T.; Vallabhaneni, S. Candida auris: The recent emergence of a multidrug-resistant fungal pathogen. Med. Mycol. 2019, 57, 1–12. 10.1093/mmy/myy054

[13] Eix, E.F.; Nett, J.E. Candida auris: Epidemiology and antifungal strategy. Annu. Rev. Med. 2025, 76, 57–67. 10.1146/annurev-med-061523-021233

[14] Maziarz, E.K.; Perfect, J.R. Cryptococcosis. Infect. Dis. Clin. North Am. 2016, 30, 179–206. 10.1016/j.idc.2015.10.006

[15] Archuleta, S.; Gharamti, A.A.; Sillau, S.; Castellanos, P.; Chadalawada, S.; Mundo, W.; Bandali, M.; Oñate, J.; Martínez, E.; Chastain, D.B.; et al. Increased mortality associated with uncontrolled diabetes mellitus in patients with pulmonary cryptococcosis: A single US cohort study. Ther. Adv. Infect. Dis. 2021, 8, 20499361211004367. 10.1177/20499361211004367

[16] Mendoza-Reyes, D.F.; Gómez-Gaviria, M.; Mora-Montes, H.M. Candida lusitaniae: Biology, pathogenicity, virulence factors, diagnosis, and treatment. Infect. Drug Resist. 2022, 15, 5121–5135. 10.2147/IDR.S383785

[17] Gharehbolagh, S.A.; Nasimi, M.; Afshari, S.A.K.; Ghasemi, Z.; Rezaie, S. First case of superficial infection due to Naganishia albida (formerly Cryptococcus albidus) in Iran: A review of the literature. Curr. Med. Mycol. 2017, 3, 33–37. 10.29252/cmm.3.2.33

[18] Oliveira, V.F.D.; Funari, A.P.; Taborda, M.; Magri, A.S.G.K.; Levin, A.S.; Magri, M.M.C. Cutaneous Naganishia albida (Cryptococcus albidus) infection: A case report and literature review. Rev. Inst. Med. Trop. Sao Paulo 2023, 65, e60. 10.1590/S1678-9946202365060

[19] Iturrieta-Gonzalez, I.A.; Padovan, A.C.B.; Bizerra, F.C.; Hahn, R.C.; Colombo, A.L. Multiple species of Trichosporon produce biofilms highly resistant to triazoles and amphotericin B. PLoS One 2014, 9, e109553. 10.1371/journal.pone.0109553

[20] Fisher, M.C.; Denning, D.W.; Cuomo, C.A. Tackling the emerging threat of antifungal resistance to human health. Science 2022, 375, eabm3442. 10.1126/science.abm3442

[21] Perfect, J.R. The antifungal pipeline: A reality check. Nat. Rev. Drug Discov. 2017, 16, 603–616. 10.1038/nrd.2017.46

[22] Faulkner, D.J. Marine natural products. Nat. Prod. Rep. 2001, 18, 1R–49R. 10.1039/B006897G

[23] Alves, C.; Silva, J.; Pinteus, S.; Gaspar, H.; Alpoim, M.C.; Botana, L.M.; Pedrosa, R. From marine origin to therapeutics: The antitumor potential of marine algae-derived compounds. Front. Pharmacol. 2018, 9, 777. 10.3389/fphar.2018.00777

[24] Mayer, A.M.S.; Rodríguez, A.D.; Taglialatela-Scafati, O.; Fusetani, N. Marine pharmacology in 2016–2017: Marine compounds with antibacterial, antidiabetic, antifungal, anti-inflammatory, antiprotozoal, antituberculosis, and antiviral activities; affecting the immune and nervous system, and other miscellaneous mechanisms of action. Mar. Drugs 2020, 18, 5. 10.3390/md18010005

[25] Carroll, A.R.; Copp, B.R.; Davis, R.A.; Keyzers, R.A.; Prinsep, M.R. Marine natural products. Nat. Prod. Rep. 2023, 40, 275–325. 10.1039/D2NP00083K

[26] Čmiková, N.; Galovičová, L.; Miškeje, M.; Borotová, P.; Kluz, M.; Kačániová, M. Determination of antioxidant, antimicrobial activity, heavy metals and elements content of seaweed extracts. Plants 2022, 11, 1493. 10.3390/plants11111493

[27] Ahmed, H.K.; Heneidak, S.; Rasmey, A.H.M.; El Shoubaky, G. In vitro comparative antimicrobial potential of bioactive crude and fatty acids extracted from abundant marine macroalgae, Egypt. Front. Sci. Res. Technol. 2023, 7. 10.21608/fsrt.2023.221539.1097

[28] Wijesekara, I.; Pangestuti, R.; Kim, S.K. Biological activities and potential health benefits of sulphated polysaccharides derived from marine algae. Carbohydr. Polym. 2011, 84, 14–21. 10.1016/j.carbpol.2010.10.062

[29] Ngo, D.H.; Kim, S.K. Sulphated polysaccharides as bioactive agents from marine algae. Int. J. Biol. Macromol. 2013, 62, 70–75. 10.1016/j.ijbiomac.2013.08.036

[30] Hentati, F.; Delattre, C.; Gardarin, C.; Desbrières, J.; Le Cerf, D.; Rihouey, C.; Michaud, P.; Abdelkafi, S.; Pierre, G. Structural features and rheological properties of a sulphated xylogalactan-rich fraction isolated from Tunisian red seaweed Jania adhaerens. Appl. Sci. 2020, 10, 1655. 10.3390/app10051655

[31] Soares, F.; Fernandes, C.; Silva, P.; Pereira, L.; Gonçalves, T. Antifungal activity of carrageenan extracts from the red alga Chondracanthus teedei var. lusitanicus. J. Appl. Phycol. 2016, 28, 2991–2998. 10.1007/s10811-016-0849-9

[32] Li, B.; Lu, F.; Wei, X.; Zhao, R. Marine natural products and their synthetic analogs as antifungal agents. Curr. Med. Chem. 2008, 15, 3430–3441. 10.2174/092986708786848523

[33] Steneck, R.S.; Dethier, M.N. A functional group approach to the structure of algal-dominated communities. Oikos 1994, 69, 476–498. 10.2307/3545860

[34] Pinedo, S.; Ballesteros, E. The role of competitor, stress-tolerant and opportunist species in the development of indexes based on rocky shore assemblages for the assessment of ecological status. Ecol. Indic. 2019, 107, 105556. 10.1016/j.ecolind.2019.105556

[35] Terradas-Fernández, M.; Valverde-Urrea, M.; Casado-Coy, N.; Sanz-Lazaro, C. The ecological condition of vermetid platforms affects the cover of the alien seaweed Caulerpa cylindracea. Sci. Mar. 2020, 84, 181–191. 10.3989/scimar.04984.06A

[36] de Azevedo, M.I.G.; Souza, P.F.N.; Monteiro Júnior, J.E.; Grangeiro, T.B. Chitosan and chitooligosaccharides: Antifungal potential and structural insights. Chem. Biodivers. 2024, 21, e202400044. 10.1002/cbdv.202400044

[37] Ke, C.L.; Deng, F.S.; Chuang, C.Y.; Lin, C.H. Antimicrobial actions and applications of chitosan. Polymers 2021, 13, 904. 10.3390/polym13060904

[38] Mármol, Z.; Páez, G.; Rincón, M.; Araujo, K.; Aiello, C.; Chandler, C.; Gutiérrez, E. Quitina y quitosano polímeros amigables. Una revisión de sus aplicaciones. Rev. Tecnocientífica URU 2013, 1, 53–58.

[39] Gildberg, A.; Stenberg, E. A new process for advanced utilisation of shrimp waste. Process Biochem. 2001, 36, 809–812. 10.1016/S0032-9592(00)00278-8

[40] Pillai, C.K.S.; Paul, W.; Sharma, C.P. Chitin and chitosan polymers: Chemistry, solubility and fiber formation. Prog. Polym. Sci. 2009, 34, 641–678. 10.1016/j.progpolymsci.2009.04.001

[41] Peña, A.; Sánchez, N.S.; Calahorra, M. Effects of chitosan on Candida albicans: Conditions for its antifungal activity. Biomed Res. Int. 2013, 2013, 527549. 10.1155/2013/527549

[42] Matica, M.A.; Aachmann, F.L.; Tøndervik, A.; Sletta, H.; Ostafe, V. Chitosan as a wound dressing starting material: Antimicrobial properties and mode of action. Int. J. Mol. Sci. 2019, 20, 5889. 10.3390/ijms20235889

[43] Lopez-Moya, F.; Colom-Valiente, M.F.; Martinez-Peinado, P.; Martinez-Lopez, J.E.; Puelles, E.; Sempere-Ortells, J.M.; Lopez-Llorca, L.V. Carbon and nitrogen limitation increase chitosan antifungal activity in Neurospora crassa and fungal human pathogens. Fungal Biol. 2015, 119, 154–169. 10.1016/j.funbio.2014.12.003

[44] Alburquenque, C.; Bucarey, S.A.; Neira-Carrillo, A.; Urzúa, B.; Hermosilla, G.; Tapia, C.V. Antifungal activity of low molecular weight chitosan against clinical isolates of Candida spp. Med. Mycol. 2010, 48, 1018–1023. 10.3109/13693786.2010.486412

[45] Palma-Guerrero, J.; Lopez-Jimenez, J.A.; Pérez-Berná, A.J.; Huang, I.C.; Jansson, H.B.; Salinas, J.; Lopez-Llorca, L.V. Membrane fluidity determines sensitivity of filamentous fungi to chitosan. Mol. Microbiol. 2010, 75, 1021–1032. 10.1111/j.1365-2958.2009.07039.x

[46] Cormaci, M.; Furnari, G.; Alongi, G. Flora marina bentonica del Mediterraneo: Rhodophyta (Rhodymeniophycidae escluse). Boll. Accad. Gioenia Sci. Nat. Catania 2017, 50, FP1–FP391. 10.35352/gioenia.v53i383.87

[47] Coat, G.; Dion, P.; Noailles, M.C.; De Reviers, B.; Fontaine, J.M.; Berger-Perrot, Y.; Loiseaux-De Goër, S. Ulva armoricana (Ulvales, Chlorophyta) from the coasts of Brittany (France). II. Nuclear rDNA ITS sequence analysis. Eur. J. Phycol. 1998, 33, 81–86. 10.1017/S0967026297001455

[48] Saunders, G.W. Applying DNA barcoding to red macroalgae: A preliminary appraisal holds promise for future applications. Philos. Trans. R. Soc. B Biol. Sci. 2005, 360, 1879–1888. 10.1098/rstb.2005.1719

[49] Hall, T.A. BioEdit: A user-friendly biological sequence alignment editor and analysis program for Windows 95/98/NT. Nucleic Acids Symp. Ser. 1999, 41, 95–98.

[50] Qi, H.; Huang, L.; Liu, X.; Liu, D.; Zhang, Q.; Liu, S. Antihyperlipidemic activity of high sulfate content derivative of polysaccharide extracted from Ulva pertusa (Chlorophyta). Carbohydr. Polym. 2012, 87, 1637–1640. 10.1016/j.carbpol.2011.09.073

[51] Yang, Q.; Jiang, Y.; Fu, S.; Shen, Z.; Zong, W.; Xia, Z.; Zang, Z.; Jiang, X. Protective effects of Ulva lactuca polysaccharide extract on oxidative stress and kidney injury induced by D-galactose in mice. Mar. Drugs 2021, 19, 539. 10.3390/md19100539

[52] AOCS. Official Methods and Recommended Practices of the American Oil Chemists’ Society 4th ed.; Method Aa 6–38; American Oil Chemists’ Society: Champaign, IL, USA, 1993.

[53] Yemm, E.W.; Willis, A.J. The estimation of carbohydrates in plant extracts by anthrone. Biochem. J. 1954, 57, 508–514. 10.1042/bj0570508

[54] Lloyd, A.G.; Dodgson, K.S.; Price, R.G.; Rose, F.A. I. Polysaccharide sulphates. Biochim. Biophys. Acta 1961, 46, 108–115. 10.1016/0006-3002(61)90652-7

[55] Bradford, M.M. A rapid and sensitive method for the quantitation of microgram quantities of protein utilizing the principle of protein-dye binding. Anal. Biochem. 1976, 72, 248–254. 10.1006/abio.1976.9999

[56] Burke, D.; Dawson, D.; Stearns, T. Methods in Yeast Genetics: A Cold Spring Harbor Laboratory Course Manual, 2000 ed.; Cold Spring Harbor Laboratory Press: Plainview, NY, USA, 2000.

[57] Elzhov, T.V.; Mullen, K.M.; Spiess, A.; Bolker, B. minpack.lm: R Interface to the Levenberg-Marquardt Nonlinear Least-Squares Algorithm Found in MINPACK, Plus Support for Bounds, R package version 1. 2–4, 2023. Available online: https://CRAN.R-project.org/package=minpack.lm

[58] R Core Team. R: A Language and Environment for Statistical Computing; R Foundation for Statistical Computing: Vienna, Austria, 2023. Available online: https://www.R-project.org/

[59] Matsuhiro, B. Vibrational spectroscopy of seaweed galactans. In Fifteenth International Seaweed Symposium; Springer: Dordrecht, The Netherlands, 1996; pp. 481–489. 10.1007/978-94-009-1659-3_69

[60] Soto-Vásquez, M.R.; Alvarado-García, P.A.A.; Youssef, F.S.; Ashour, M.L.; Bogari, H.A.; Elhady, S.S. FTIR characterization of sulphated polysaccharides obtained from Macrocystis integrifolia algae and verification of their antiangiogenic and immunomodulatory potency in vitro and in vivo. Mar. Drugs 2023, 21, 36. 10.3390/md21010036

[61] Gómez-Ordóñez, E.; Rupérez, P. FTIR-ATR spectroscopy as a tool for polysaccharide identification in edible brown and red seaweeds. Food Hydrocoll. 2011, 25, 1514–1520. 10.1016/j.foodhyd.2011.02.009

[62] Sfriso, A.A.; Gallo, M.; Baldi, F. Seasonal variation and yield of sulphated polysaccharides in seaweeds from the Venice Lagoon. Bot. Mar. 2017, 60, 339–349. 10.1515/bot-2016-0063

[63] Wang, Z.; Sun, Q.; Zhang, H.; Wang, J.; Fu, Q.; Qiao, H.; Wang, Q. Insight into antibacterial mechanism of polysaccharides: A review. LWT 2021, 150, 111929. 10.1016/j.lwt.2021.111929

[64] Afzal, S.; Yadav, A.K.; Poonia, A.K.; Choure, K.; Yadav, A.N.; Pandey, A. Antimicrobial therapeutics isolated from algal source: Retrospect and prospect. Biologia 2023, 78, 291–305. 10.1007/s11756-022-01207-3

[65] Vishwakarma, J.; Vavilala, S.L. Evaluating the antibacterial and antibiofilm potential of sulphated polysaccharides extracted from green algae Chlamydomonas reinhardtii. J. Appl. Microbiol. 2019, 127, 1004–1017. 10.1111/jam.14364

[66] Jaime, M.D.L.A.; Lopez-Llorca, L.V.; Conesa, A.; Lee, A.Y.; Proctor, M.; Heisler, L.E.; Gebbia, M.; Giaever, G.; Westwood, J.T.; Nislow, C. Identification of yeast genes that confer resistance to chitosan oligosaccharide (COS) using chemogenomics. BMC Genomics 2012, 13, 267. 10.1186/1471-2164-13-267

[67] Lopez-Moya, F.; Suarez-Fernandez, M.; Lopez-Llorca, L.V. Molecular mechanisms of chitosan interactions with fungi and plants. Int. J. Mol. Sci. 2019, 20, 332. 10.3390/ijms20020332

[68] Shih, P.Y.; Liao, Y.T.; Tseng, Y.K.; Deng, F.S.; Lin, C.H. A potential antifungal effect of chitosan against Candida albicans is mediated via the inhibition of SAGA complex component expression and the subsequent alteration of cell surface integrity. Front. Microbiol. 2019, 10, 602. 10.3389/fmicb.2019.00602

[69] Lopez-Moya, F., Martin-Urdiroz, M., Oses-Ruiz, M., Were, V. M., Fricker, M. D., Littlejohn, G., Lopez-Llorca, L.V. & Talbot, N. J. Chitosan inhibits septin-mediated plant infection by the rice blast fungus Magnaporthe oryzae in a protein kinase C and Nox1 NADPH oxidase-dependent manner. New Phytologist, 2021, 230(4), 1578–1593.

[70] Guo, Z.; Xing, R.; Liu, S.; Zhong, Z.; Ji, X.; Wang, L.; Li, P. The influence of molecular weight of quaternized chitosan on antifungal activity. Carbohydr. Polym. 2008, 71, 694–697. 10.1016/j.carbpol.2007.06.027

[71] Bagheri-Khoulenjani, S.; Taghizadeh, S.M.; Mirzadeh, H. An investigation on the short-term biodegradability of chitosan with various molecular weights and degrees of deacetylation. Carbohydr. Polym. 2009, 78, 773–778. 10.1016/j.carbpol.2009.06.020

[72] Aranda-Martinez, A.; Lopez-Moya, F.; Lopez-Llorca, L.V. Cell wall composition plays a key role on sensitivity of filamentous fungi to chitosan. J. Basic Microbiol. 2016, 56, 1059–1070. 10.1002/jobm.201500775

[73] Mishra, P.; Prasad, R. An overview of lipids of Candida albicans. Prog. Lipid Res. 1990, 29, 65–85. 10.1016/0163-7827(90)90006-7

[74] Chowdhary, A.; Sharma, C.; Meis, J.F. Candida auris: A rapidly emerging cause of hospital-acquired multidrug-resistant fungal infections globally. PLoS Pathog. 2017, 13, e1006290. 10.1371/journal.ppat.1006290

[75] Pallotta, F.; Viale, P.; Barchiesi, F. Candida auris: The new fungal threat. Le Infez. Med. 2023, 31, 323–330. 10.53854/liim-3103-6

[76] Yen, M.T.; Yang, J.H.; Mau, J.L. Antioxidant properties of chitosan from crab shells. Carbohydr. Polym. 2008, 74, 840–844. 10.1016/j.carbpol.2008.05.003

