## Supplementary figures and tables for "Species-dependent antifungal profiles reveal stronger yeast inhibition by chitosan than by a sulfate-containing polysaccharide-rich extract from *Jania adhaerens*"

Supplementary material

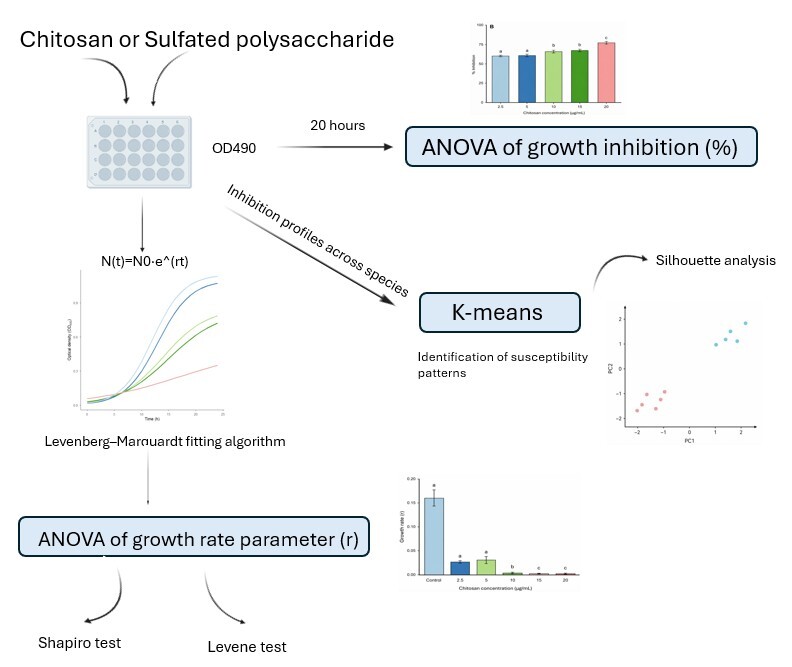

**Figure S1.** Schematic representation of the statistical workflow applied to evaluate the antifungal activity of chitosan and sulfated polysaccharides against yeast species. Growth kinetics were monitored over 24 h using OD490 measurements and fitted to an exponential growth model using the Levenberg–Marquardt algorithm to obtain the growth rate parameter (r), which was subsequently analyzed by ANOVA. In parallel, growth inhibition percentages at 20 h were calculated and analyzed by ANOVA. Species susceptibility patterns were further explored through K-means clustering based on inhibition profiles across concentrations, with the optimal number of clusters determined using silhouette analysis.

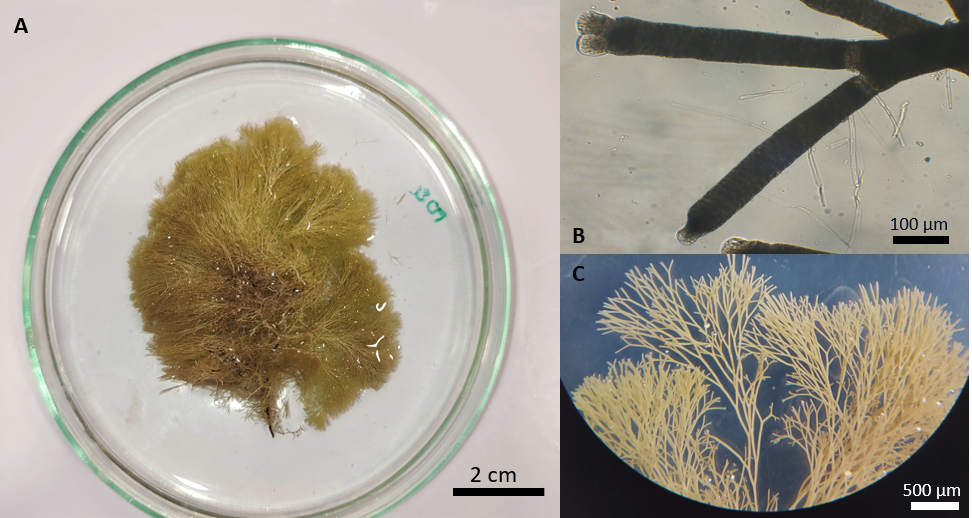

**Figure S2.** Morphological characterization of *Jania adhaerens*. (A) General appearance of the alga, showing a dichotomous thallus with a pale pink-lilac coloration and an approximate length of 5 cm. (B) Detail of the calcified intergenicula, which presented diameters ranging from 80 to 120 µm and lengths approximately 5–6 times greater than their width. (C) Representative branching pattern of the thallus, with dichotomy angles greater than 30°.

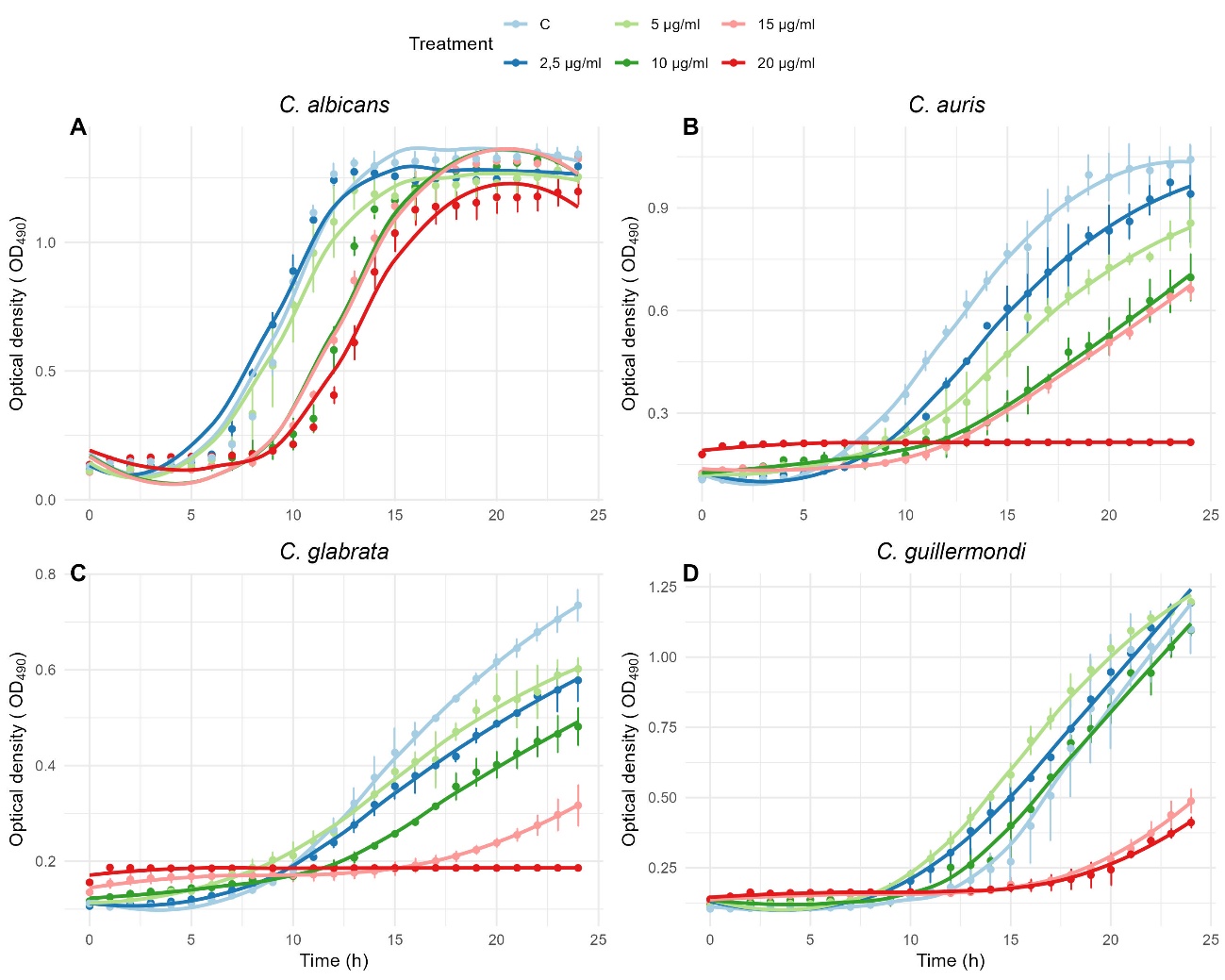

Figure S3: Growth kinetics of *C. albicans* (A), *C. auris* (B), *Cr. bacillisporus* (C) and *Cr. deuterogatti* (D) exposed to different concentrations of chitosan over 24 h. Growth was monitored by OD490 measurements. Treatment indicates the chitosan concentrations applied. C: control. Values represent mean ± standard deviation.

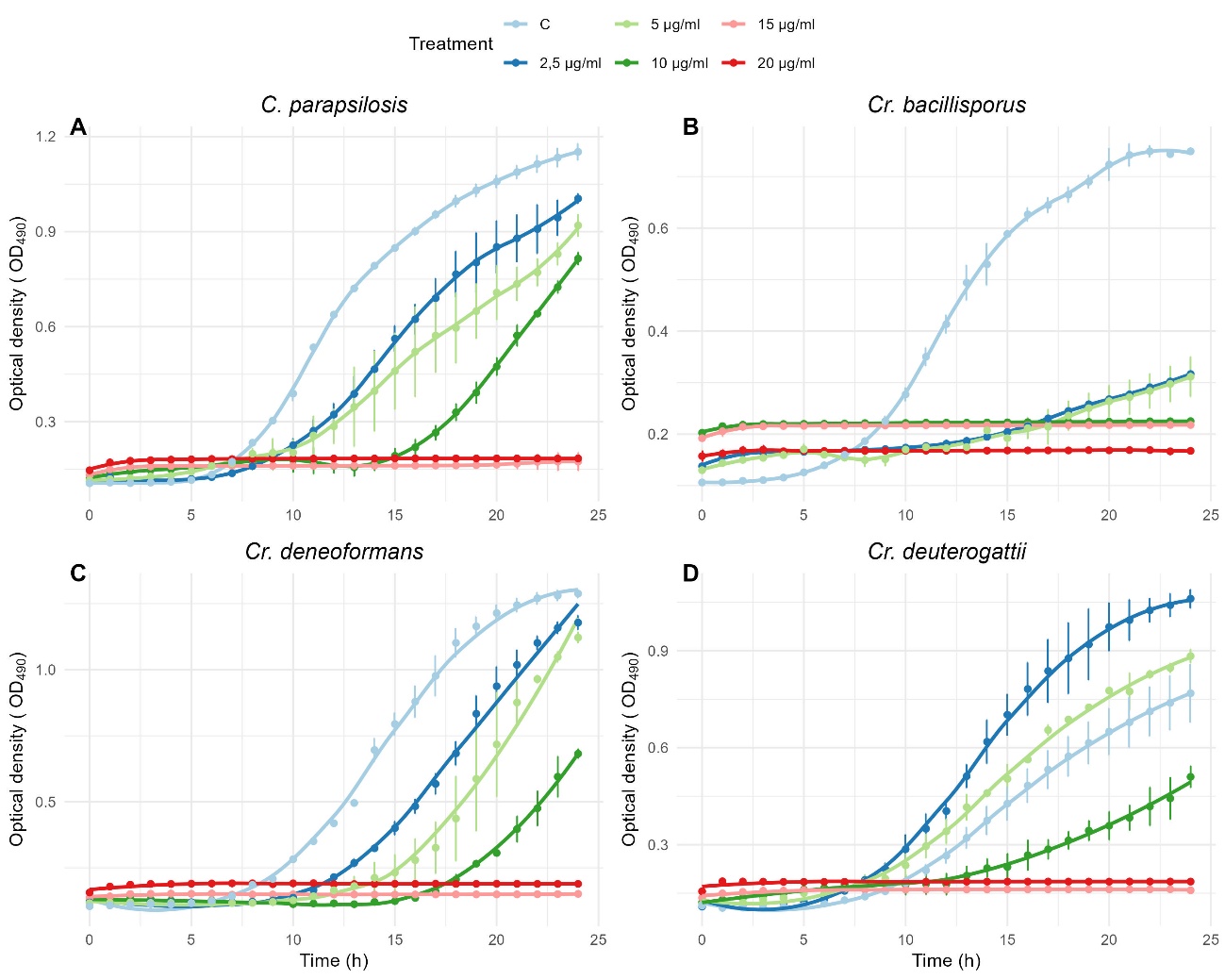

Figure S4: Growth kinetics of *C. parapsilosis* (A) and *Cryptococcus* spp. including *Cr. bacillisporus* (B), *Cr. deneoformans* (C) and *Cr. deuterogatti* (D) exposed to different concentrations of chitosan over 24 h. Growth was monitored by OD490 measurements. Treatment indicates the chitosan concentrations applied. C: control. Values represent mean ± standard deviation.

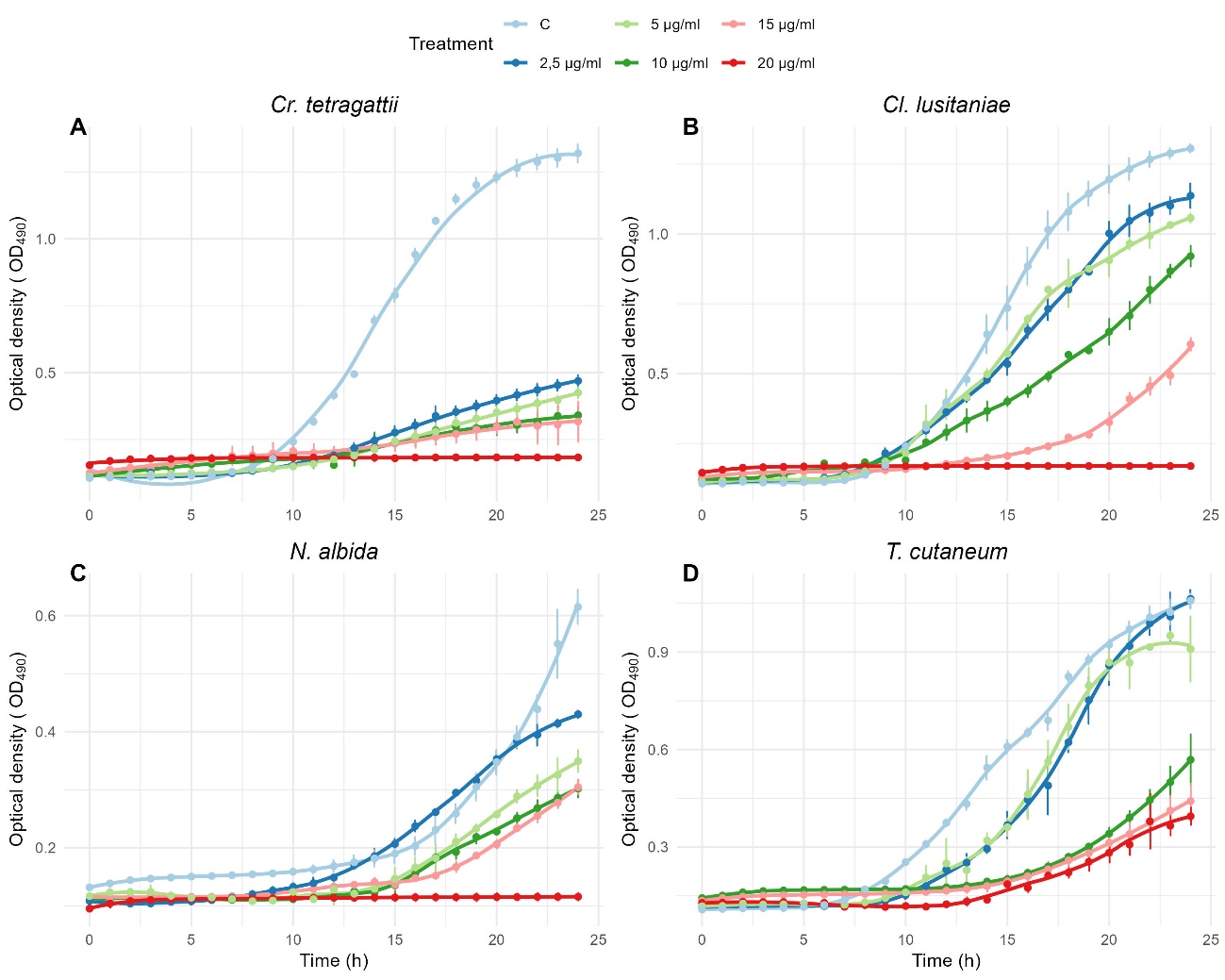

Figure S5: Growth kinetics of *Cr. tetragattii* (A), *Cl. lusitaniae* (B), *N. albida* (C) and *T. cutaneum* (D) exposed to different concentrations of chitosan over 24 h. Growth was monitored by OD490 measurements. Treatment indicates the chitosan concentrations applied. C: control. Values represent mean ± standard deviation.

Table S1: Growth rate parameter (r) obtained from the exponential growth model fitted for each yeast species exposed to different concentrations of chitosan. Estimation values, standard errors, t values and p values are shown for each treatment condition.

| Species | Treatment | Estimate | Standard error | T value | p.value |
| --- | --- | --- | --- | --- | --- |
| *C. albicans* |  |  |  |  |  |
|  | C | 0.186 | 0.003 | 58.04 | **<0.001** |
|  | 2.5 µg/ml | 0.185 | 0.003 | 56.93 | **<0.001** |
|  | 5 µg/ml | 0.178 | 0.003 | 55.72 | **<0.001** |
|  | 10 µg/ml | 0.159 | 0.003 | 40.23 | **<0.001** |
|  | 15 µg/ml | 0.156 | 0.003 | 45.64 | **<0.001** |
|  | 20 µg/ml | 0.141 | 0.004 | 34.8 | **<0.001** |
| *C. auris* |  |  |  |  |  |
|  | C | 0.429 | 0.020 | 20.82 | **<0.001** |
|  | 2.5 µg/ml | 0.363 | 0.019 | 18.84 | **<0.001** |
|  | 5 µg/ml | 0.394 | 0.044 | 8.85 | **<0.001** |
|  | 10 µg/ml | 0.148 | 0.032 | 4.562 | **<0.001** |
|  | 15 µg/ml | 0.204 | 0.023 | 8.61 | **<0.001** |
|  | 20 µg/ml | 0.050 | 0.006 | 7.62 | **<0.001** |
| *C. glabrata* |  |  |  |  |  |
|  | C | 0.114 | 0.004 | 28.32 | **<0.001** |
|  | 2.5 µg/ml | 0.094 | 0.002 | 32.75 | **<0.001** |
|  | 5 µg/ml | 0.090 | 0.003 | 29.47 | **<0.001** |
|  | 10 µg/ml | 0.081 | 0.002 | 29.10 | **<0.001** |
|  | 15 µg/ml | 0.022 | 0.002 | 9.08 | **<0.001** |
|  | 20 µg/ml | 0.0002 | 0.0001 | 2.70 | **<0.001** |
| *C. guillermondii* |  |  |  |  |  |
|  | C | 0.125 | 0.027 | 4.530 | **<0.001** |
|  | 2.5 µg/ml | 0.175 | 0.023 | 7.631 | **<0.001** |
|  | 5 µg/ml | 0.181 | 0.010 | 71.008 | **<0.001** |
|  | 10 µg/ml | 0.189 | 0.037 | 5.019 | **<0.001** |
|  | 15 µg/ml | 0.036 | 0.009 | 3.943 | **0.001** |
|  | 20 µg/ml | 0.017 | 0.006 | 2.71 | **0.015** |
| *C. parapsilosis* |  |  |  |  |  |
|  | C | 0.173 | 0.009 | 18.9 | **<0.001** |
|  | 2.5 µg/ml | 0.177 | 0.007 | 23.03 | **<0.001** |
|  | 5 µg/ml | 0.130 | 0.017 | 7.49 | **<0.001** |
|  | 10 µg/ml | 0.003 | 0.009 | 0.417 | 0.63 |
|  | 15 µg/ml | 0.001 | 0.0001 | 5.90 | **<0.001** |
| *Cr. bacillisporus* |  |  |  |  |  |
|  | C | 0.158 | 0.005 | 29.24 | **<0.001** |
|  | 2.5 µg/ml | 0.020 | 0.001 | 15.85 | **<0.001** |
|  | 5 µg/ml | 0.023 | 0.004 | 5.037 | **<0.001** |
|  | 10 µg/ml | 0.001 | 0.001 | 1.318 | 0.197 |
|  | 15 µg/ml | 0.0004 | 0.0006 | 0.775 | 0.444 |
|  | 20 µg/ml | 0.0004 | 0.0004 | 1.016 | 0.317 |
| *Cr. deneoformans* |  |  |  |  |  |
|  | C | 0.168 | 0.005 | 29.79 | **<0.001** |
|  | 2.5 µg/ml | 0.176 | 0.004 | 40.21 | **<0.001** |
|  | 5 µg/ml | 0.177 | 0.155 | 11.409 | **<0.001** |
|  | 10 µg/ml | 0.076 | 0.008 | 9.475 | **<0.001** |
|  | 15 µg/ml | -0.006 | 0.002 | -0.322 | 0.749 |
|  | 20 µg/ml | -0.0006 | 0.0005 | -1.23 | 0.226 |
| *Cr. deuterogatti* |  |  |  |  |  |
|  | C | 0.166 | 0.014 | 11.37 | **<0.001** |
|  | 2.5 µg/ml | 0.180 | 0.014 | 12.835 | **<0.001** |
|  | 5 µg/ml | 0.142 | 0.013 | 10.44 | **<0.001** |
|  | 10 µg/ml | 0.065 | 0.021 | 3.098 | **<0.01** |
|  | 15 µg/ml | 0.0008 | 0.007 | 0.117 | 0.908 |
|  | 20 µg/ml | 0.0004 | 0.0005 | 0.863 | 0.401 |
| *Cr. tetragattii* |  |  |  |  |  |
|  | C | 0.131 | 0.008 | 16.003 | **<0.001** |
|  | 2.5 µg/ml | 0.091 | 0.004 | 18.33 | **<0.001** |
|  | 5 µg/ml | 0.087 | 0.004 | 18.09 | **<0.001** |
|  | 10 µg/ml | 0.064 | 0.007 | 8.375 | **<0.001** |
|  | 15 µg/ml | 0.045 | 0.006 | 6.662 | **<0.001** |
| *Cl. lusitaniae* |  |  |  |  |  |
|  | C | 0.177 | 0.001 | 152.7 | **<0.001** |
|  | 2.5 µg/ml | 0.159 | 0.0007 | 211.3 | **<0.001** |
|  | 5 µg/ml | 0.163 | 0.0009 | 181.2 | **<0.001** |
|  | 10 µg/ml | 0.137 | 0.001 | 103.7 | **<0.001** |
|  | 15 µg/ml | 0.096 | 0.001 | 70.26 | **<0.001** |
|  | 20 µg/ml | 0.084 | 0.002 | 29.39 | 0.226 |
| *N. albida* |  |  |  |  |  |
|  | C | 0.100 | 0.001 | 95.67 | **<0.001** |
|  | 2.5 µg/ml | 0.093 | 0.006 | 149.7 | **<0.001** |
|  | 5 µg/ml | 0.081 | 0.0006 | 122.8 | **<0.001** |
|  | 10 µg/ml | 0.075 | 0.0005 | 146.8 | **<0.001** |
|  | 15 µg/ml | 0.072 | 0.0006 | 116.2 | **<0.001** |
|  | 20 µg/ml | 0.040 | 0.001 | 35.03 | **<0.001** |
| *T. cutaneum* |  |  |  |  |  |
|  | C | 0.121 | 0.004 | 26.27 | **<0.001** |
|  | 2.5 µg/ml | 0.172 | 0.005 | 30.625 | **<0.001** |
|  | 5 µg/ml | 0.170 | 0.006 | 26.00 | **<0.001** |
|  | 10 µg/ml | 0.066 | 0.003 | 17.16 | **<0.001** |
|  | 15 µg/ml | 0.066 | 0.004 | 14.59 | **<0.001** |
|  | 20 µg/ml | 0.088 | 0.005 | 17.07 | **<0.001** |

Table S2: One-way ANOVA results for the growth rate parameter (r) of the different yeast species exposed to different concentrations of chitosan. Df: degrees of freedom. Sum Sq: sum of squares. Mean Sq: mean squares. F value: F statistic value.

| Species | Df | Sum Sq | Mean Sq | F value | p.value |
| --- | --- | --- | --- | --- | --- |
| *C. albicans* | 5 | 0.351 | 0.070 | 123.4 | **<0.001** |
|  | 12 | 0.006 | 0.0005 |  |  |
| *C. auris* | 5 | 0.066 | 0.013 | 23.28 | **<0.001** |
|  | 12 | 0.006 | 0.0005 |  |  |
| *C. glabrata* | 5 | 0.058 | 0.011 | 128.7 | **<0.001** |
|  | 12 | 0.001 | 0.00009 |  |  |
| *C. guillermondii* | 5 | 0.086 | 0.017 | 19.93 | **<0.001** |
|  | 12 | 0.104 | 0.0008 |  |  |
| *C. parapsilosis* | 5 | 0.114 | 0.022 | 117.4 | **<0.001** |
|  | 12 | 0.002 | 0.0001 |  |  |
| *Cr. bacillisporus* | 5 | 0.057 | 0.011 | 519.8 | **<0.001** |
|  | 12 | 0.0006 | 0.00002 |  |  |
| *Cr. deneoformans* | 5 | 0.132 | 0.026 | 91.52 | **<0.001** |
|  | 12 | 0.003 | 0.0002 |  |  |
| *C. deuterogatti* | 5 | 0.100 | 0.020 | 45.28 | **<0.001** |
|  | 12 | 0.005 | 0.0004 |  |  |
| *C. tetragattii* | 5 | 0.029 | 0.005 | 44.45 | **<0.001** |
|  | 12 | 0.001 | 0.001 |  |  |
| *Cl. lusitaniae* | 5 | 0.020 | 0.004 | 1033 | **<0.001** |
|  | 12 | 0.00004 | 0.00004 |  |  |
| *N. albida* | 5 | 0.005 | 0.001 | 206.4 | **<0.001** |
|  | 12 | 0.00006 | 0.000005 |  |  |
| *T. cutaneum* | 5 | 0.014 | <0.001 | 153 | **<0.001** |
|  | 12 | 0.0002 | <0.001 |  |  |

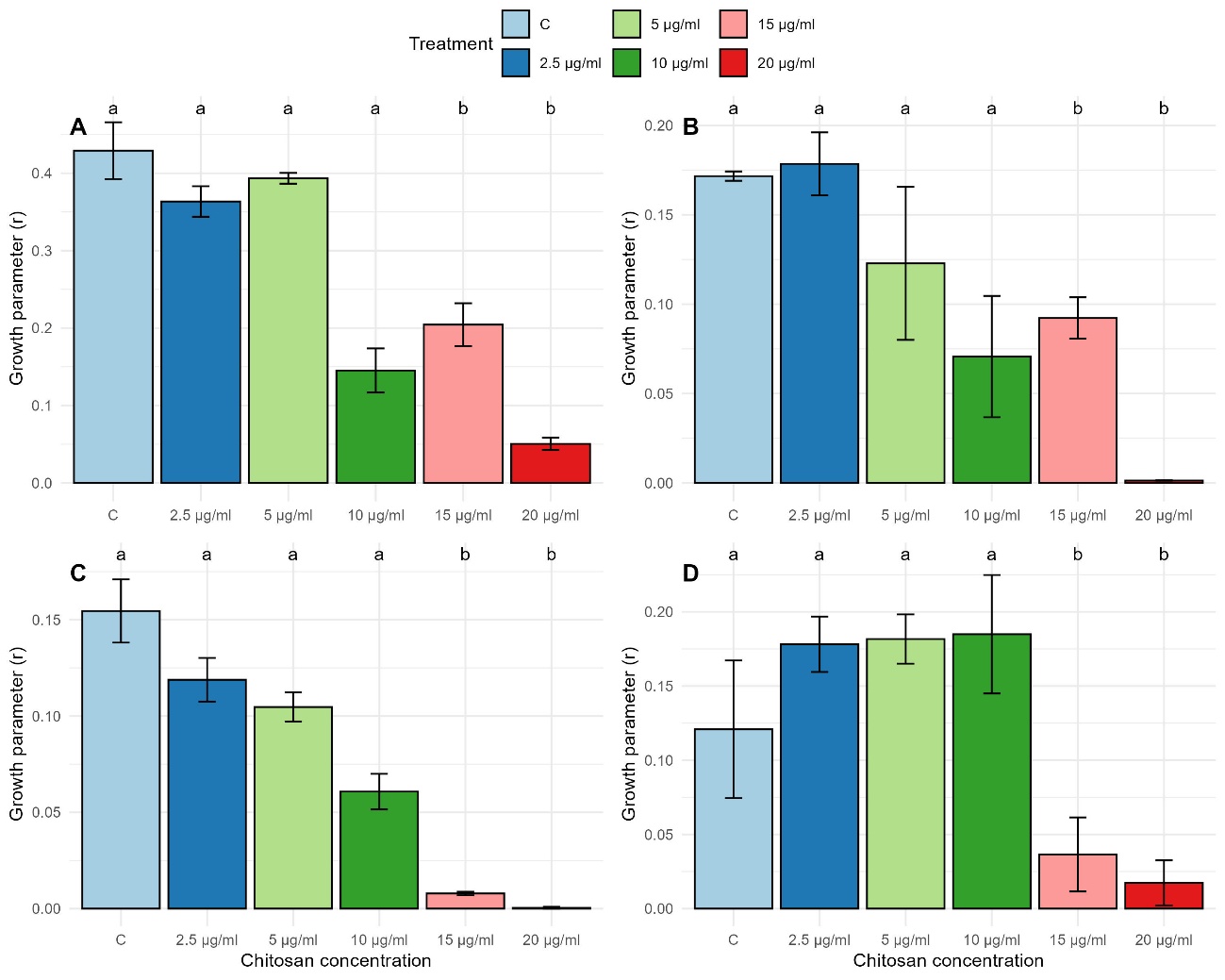

Figure S6. Effect of chitosan on the growth parameter (r) of A) *C. albicans*, B) *C. auris*, C) *C. glabrata*, and D) *C. guilliermondii* at different concentrations (2.5, 5, 10, 15, and 20 µg mL⁻¹). Bars represent the mean ± standard deviation. Different letters indicate significant differences among treatments according to Tukey’s post hoc test (p < 0.05).

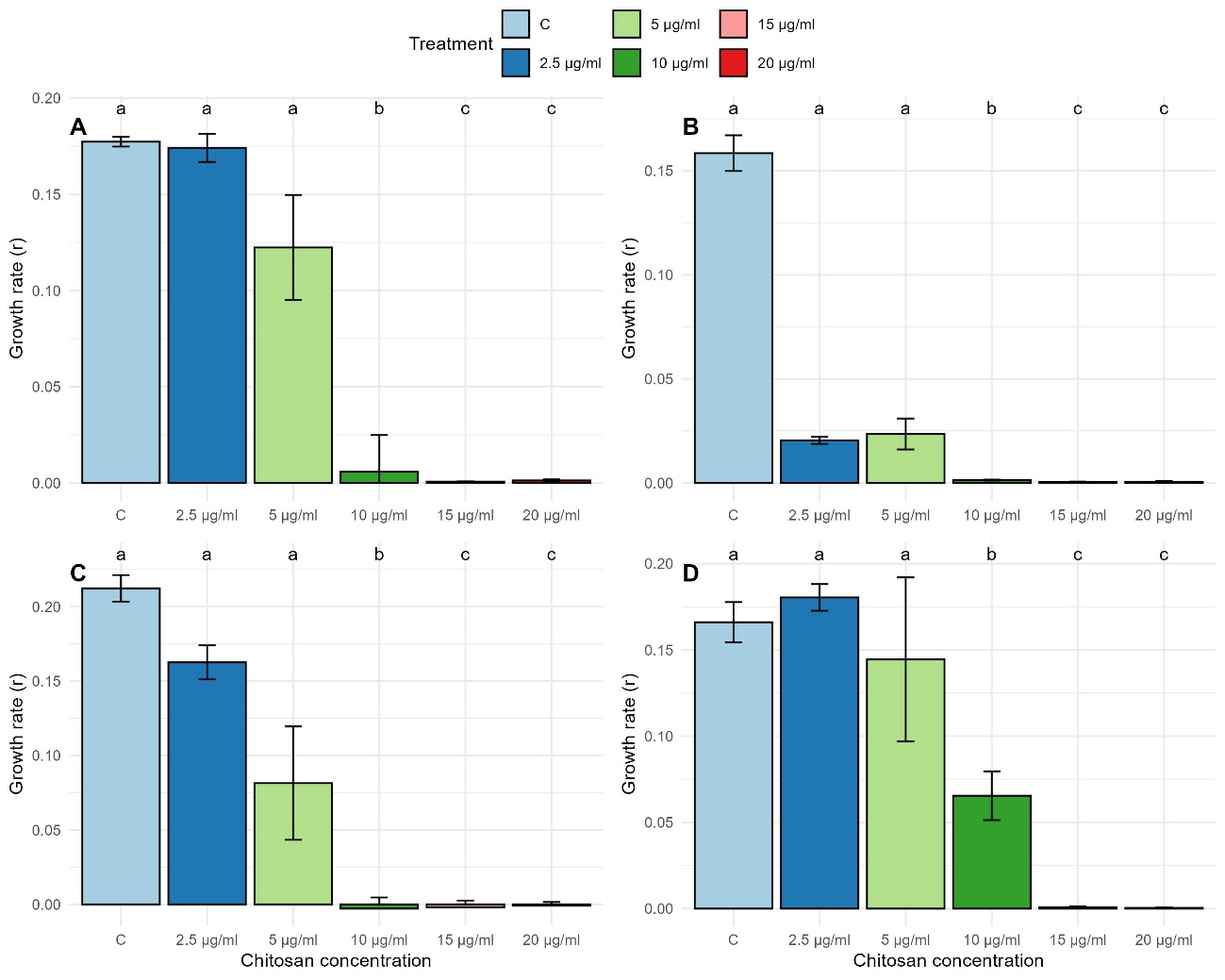

Figure S7. Effect of chitosan on the growth parameter (r) of A) *C. parapsilosis*, B) *Cr. bacillisporus*, C) *Cr. deneoformans* and D) *Cr. deuterogatti* at different concentrations (2.5, 5, 10, 15, and 20 µg mL⁻¹). Bars represent the mean ± standard deviation. Different letters indicate significant differences among treatments according to Tukey’s post hoc test (p < 0.05).

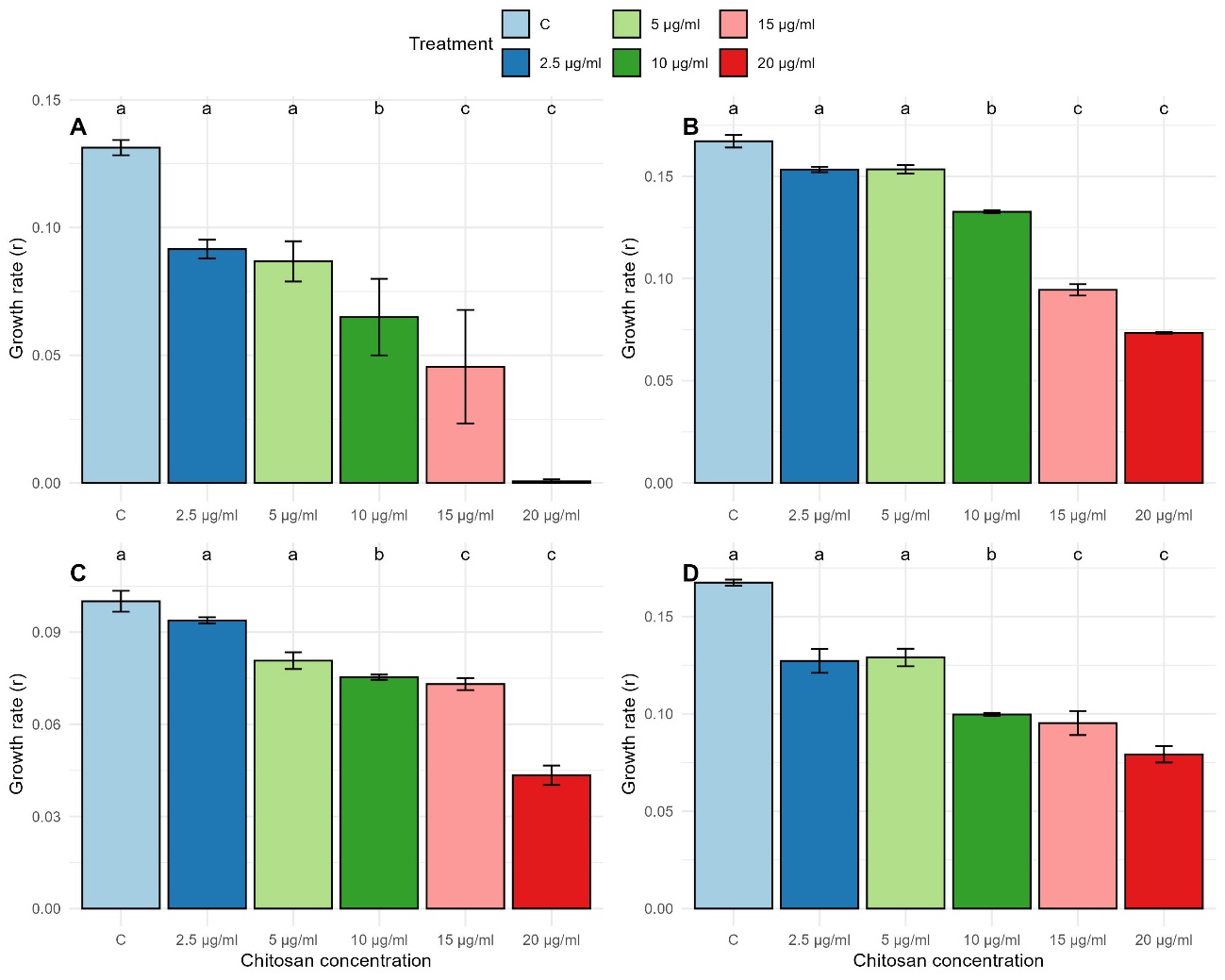

Figure S8. Effect of chitosan on the growth parameter (r) of A) *Cr. tetragattii*, B) *Cl. lusitaniae*, C) *N. albida* y D) *T. cutaneum* at different concentrations (2.5, 5, 10, 15, and 20 µg mL⁻¹). Bars represent the mean ± standard deviation. Different letters indicate significant differences among treatments according to Tukey’s post hoc test (p < 0.05).

Table S3: One-way ANOVA results for the growth inhibition of the different yeast species exposed to different concentrations of chitosan. Df: degrees of freedom. Sum Sq: sum of squares. Mean Sq: mean squares. F value: F statistic value.

| Species | Df | Sum Sq | Mean Sq | F value | p.value |
| --- | --- | --- | --- | --- | --- |
| *C. albicans* | 4 | 211 | 52.75 | 4.12 | **<0.001** |
|  | 10 | 128 | 12.80 |  |  |
| *C. auris* | 4 | 6925 | 1731 | 75.22 | **<0.001** |
|  | 10 | 230 | 23 |  |  |
| *C. glabrata* | 4 | 7465 | 1866.3 | 117.8 | **<0.001** |
|  | 10 | 158 | 15.8 |  |  |
| *C. guillermondii* | 4 | 23950 | 5987 | 14.48 | **<0.001** |
|  | 10 | 4135 | 414 |  |  |
| *C. parapsilosis* | 4 | 10118 | 2529.5 | 83.08 | **<0.001** |
|  | 10 | 304 | 30.4 |  |  |
| *Cr. bacillisporus* | 4 | 372.6 | 93.16 | 31.57 | **<0.001** |
|  | 10 | 29.5 | 2.95 |  |  |
| *Cr. deneoformans* | 4 | 9954 | 2488.4 | 34.01 | **<0.001** |
|  | 10 | 732 | 73.2 |  |  |
| *C. deuterogatti* | 4 | 38309 | 9577 | 82.03 | **<0.001** |
|  | 10 | 1168 | 117 |  |  |
| *C. tetragattii* | 4 | 503.0 | 125.75 | 38.27 | **<0.001** |
|  | 10 | 32.9 | 3.9 |  |  |
| *Cl. lusitaniae* | 4 | 11028 | 2765.9 | 428.2 | **<0.001** |
|  | 10 | 64 | 6.4 |  |  |
| *N. albida* | 4 | 4243 | 1061 | 96.43 | **<0.001** |
|  | 10 | 110 | 11 |  |  |
| *T. cutaneum* | 4 | 12933 | 3233 | 131.9 | **<0.001** |
|  | 10 | 245 | 25 |  |  |

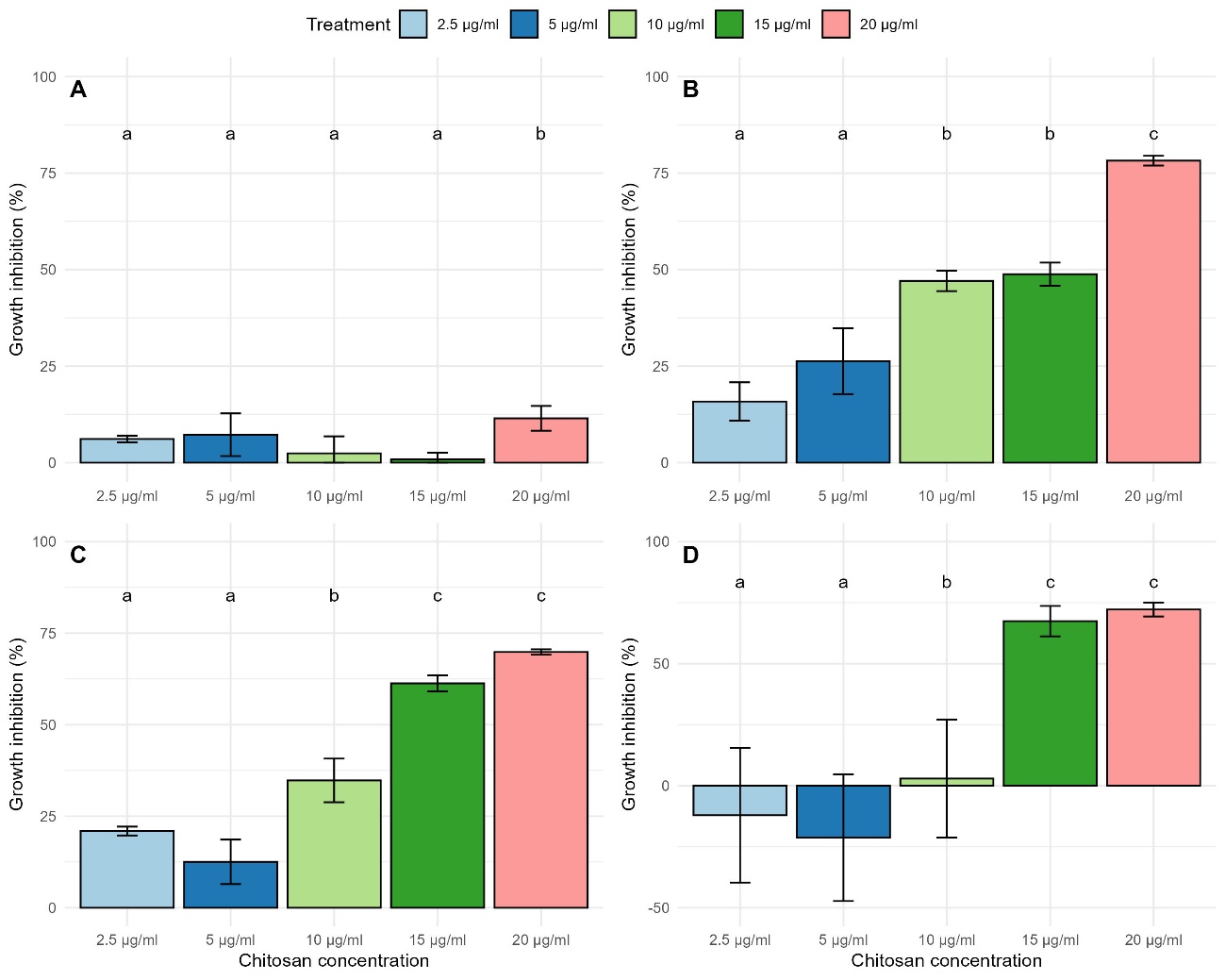

Figure S9. Effect of chitosan on growth inhibition (%) of (A) *C. albicans*, (B) *C. auris*, (C) *C. glabrata*, and (D) *C. guilliermondii* at different concentrations (2.5, 5, 10, 15, and 20 µg mL⁻¹). Bars represent the mean ± standard deviation. Different letters indicate significant differences among treatments according to Tukey’s post hoc test (p < 0.05).

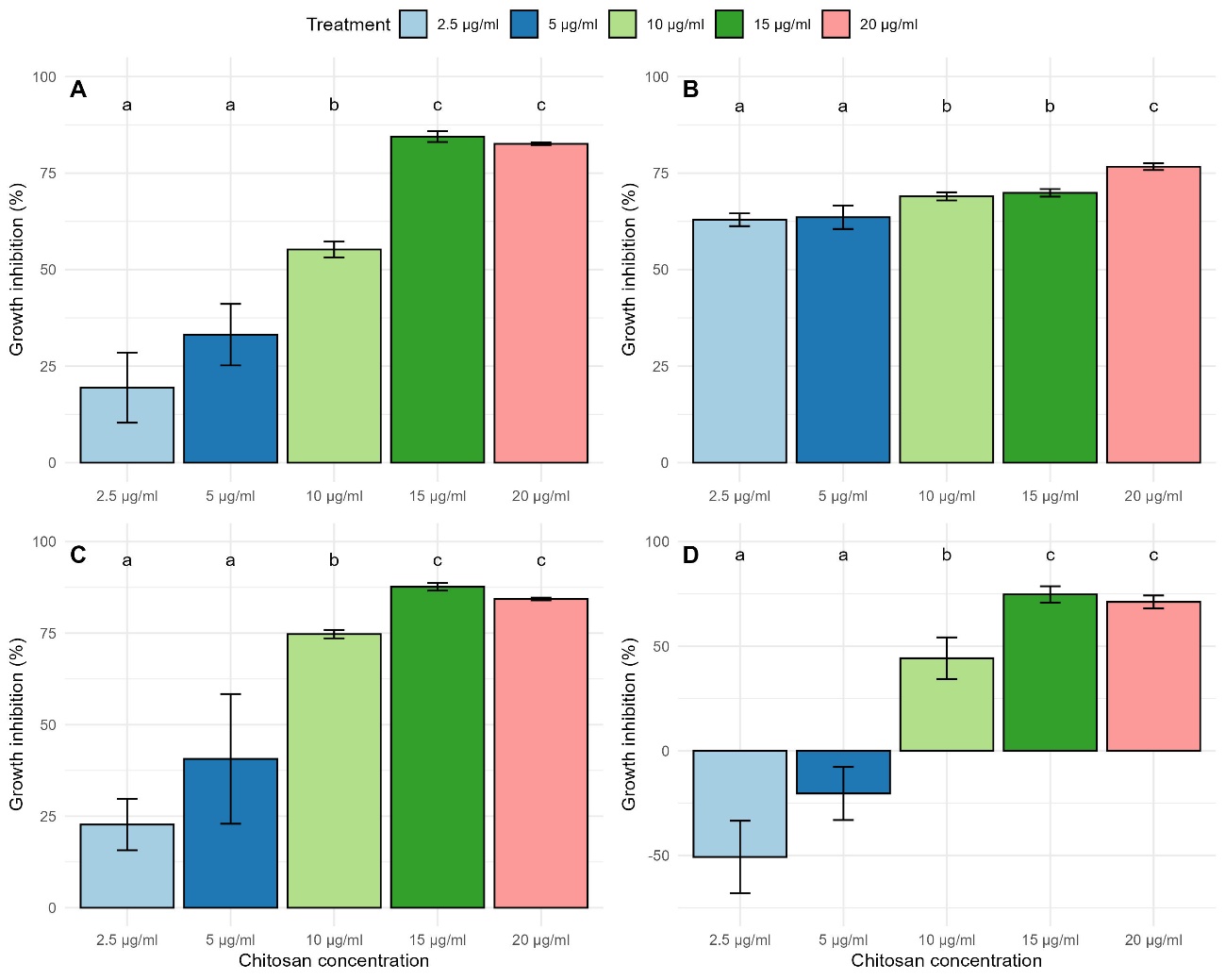

Figure S10. Effect of chitosan on growth inhibition (%) of A) *C. parapsilosis*, B) *Cr. bacillisporus*, C) *Cr. deneoformans* and D) *Cr. deuterogatti* at different concentrations (2.5, 5, 10, 15, and 20 µg mL⁻¹). Bars represent the mean ± standard deviation. Different letters indicate significant differences among treatments according to Tukey’s post hoc test (p < 0.05).

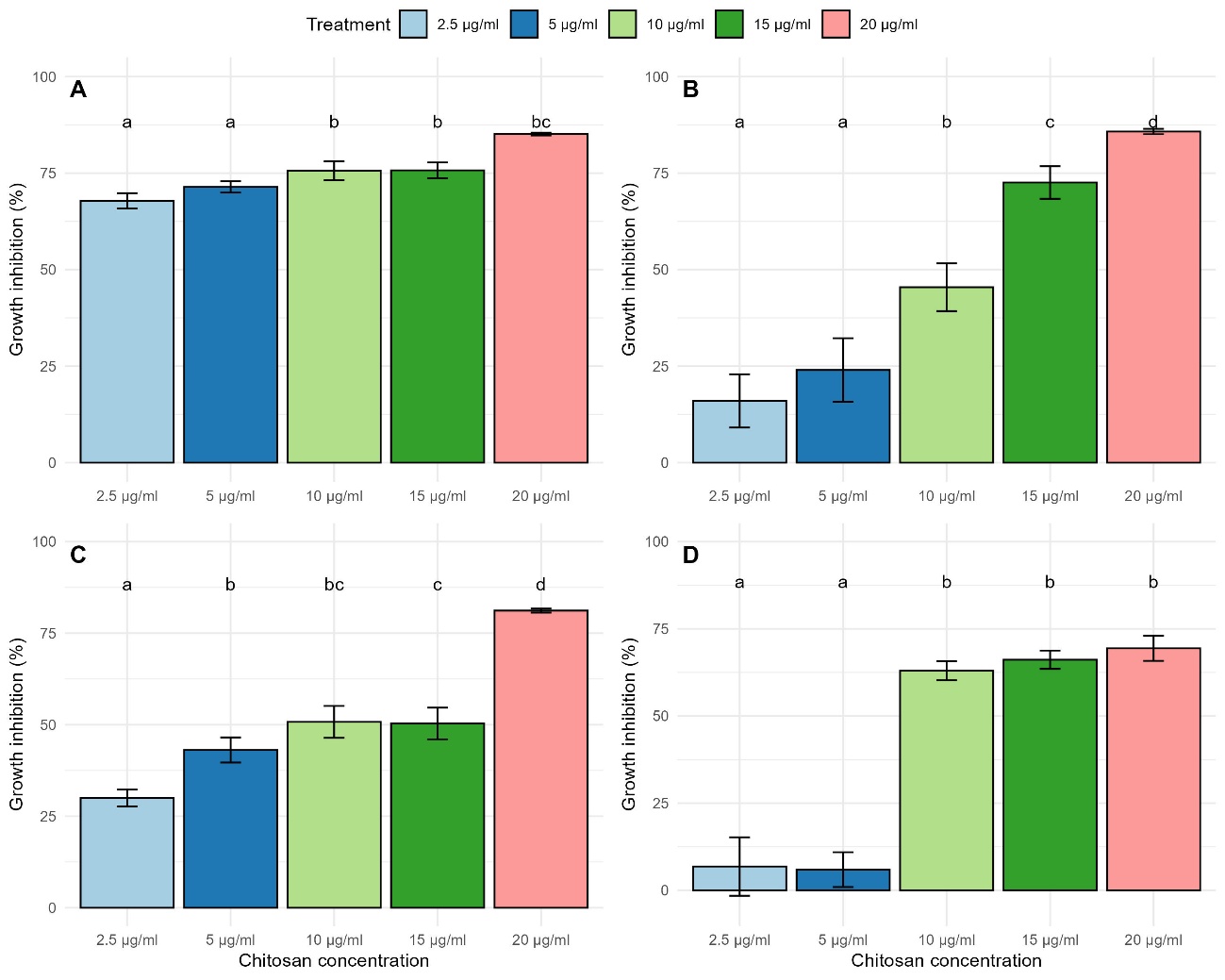

Figure S11. Effect of chitosan on growth inhibition (%) of A) *Cr. tetragattii*, B) *Cl. lusitaniae*, C) *N. albida* and D) *T. cutaneum* at different concentrations (2.5, 5, 10, 15, and 20 µg mL⁻¹). Bars represent the mean ± standard deviation. Different letters indicate significant differences among treatments according to Tukey’s post hoc test (p < 0.05).

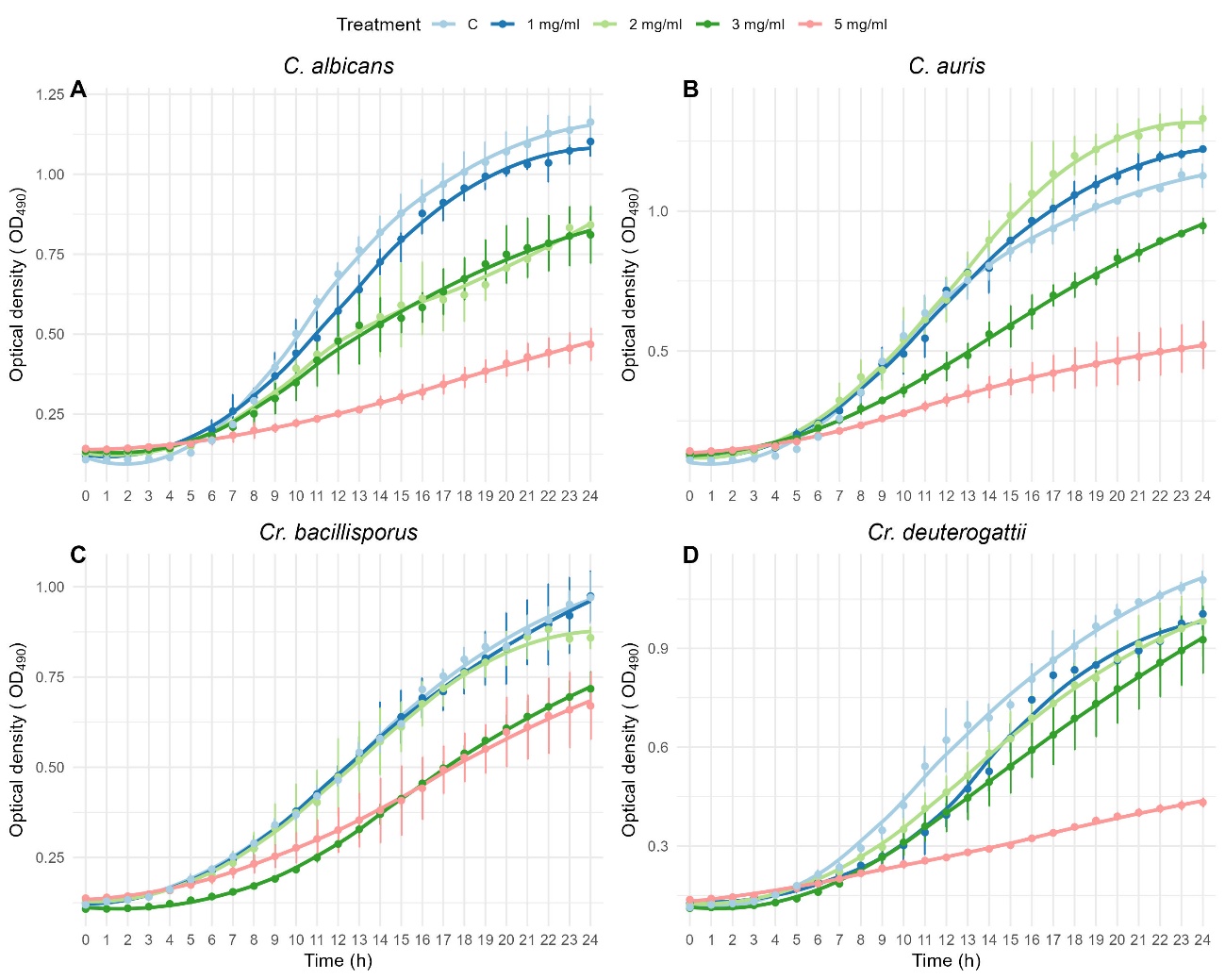

Figure S12: Growth kinetics of *C. albicans* (A), *C. auris* (B), *Cr. bacillisporus* (C) and *Cr. deuterogattii* (D) exposed to different concentrations of seaweed polysaccharides over 24 h. Growth was monitored by OD490 measurements. Treatment indicates the polysaccharides concentrations applied. C: control. Values represent mean ± standard deviation.

Table S4: Growth rate parameter (r) obtained from the exponential growth model fitted for each yeast species exposed to different concentrations of seaweed polysaccharides. Estimation values, standard errors, t values and p values are shown for each treatment condition.

| Species | Treatment | Estimate | Standard error | T value | p.value |
| --- | --- | --- | --- | --- | --- |
| *C. albicans* |  |  |  |  |  |
|  | C | 0.194 | 0.001 | 121.4 | **<0.001** |
|  | 1 mg mL⁻¹ | 0.184 | 0.002 | 90.25 | **<0.001** |
|  | 2 mg mL⁻¹ | 0.169 | 0.002 | 64.07 | **<0.001** |
|  | 3 mg mL⁻¹ | 0.167 | 0.002 | 64.75 | **<0.001** |
|  | 5 mg mL⁻¹ | 0.125 | 0.002 | 44.59 | **<0.001** |
| *C. auris* |  |  |  |  |  |
|  | C | 0.203 | 0.002 | 89.02 | **<0.001** |
|  | 1 mg mL⁻¹ | 0.202 | 0.001 | 103.6 | **<0.001** |
|  | 2 mg mL⁻¹ | 0.203 | 0.002 | 76.37 | **<0.001** |
|  | 3 mg mL⁻¹ | 0.171 | 0.002 | 67.15 | **<0.001** |
|  | 5 mg mL⁻¹ | 0.148 | 0.002 | 52.8 | **<0.001** |
| *Cr. bacillisporus* |  |  |  |  |  |
|  | C | 0.116 | 0.0005 | 195.1 | **<0.001** |
|  | 1 mg mL⁻¹ | 0.116 | 0.001 | 106.5 | **<0.001** |
|  | 2 mg mL⁻¹ | 0.114 | 0.001 | 67.21 | **<0.001** |
|  | 3 mg mL⁻¹ | 0.084 | 0.001 | 56.25 | **<0.001** |
|  | 5 mg mL⁻¹ | 0.089 | 0.001 | 45.8 | **<0.001** |
| *Cr. deuterogatti* |  |  |  |  |  |
|  | C | 0.172 | 0.002 | 65.71 | **<0.001** |
|  | 1 mg mL⁻¹ | 0.158 | 0.001 | 102.5 | **<0.001** |
|  | 2 mg mL⁻¹ | 0.160 | 0.002 | 78.38 | **<0.001** |
|  | 3 mg mL⁻¹ | 0.151 | 0.002 | 71.38 | **<0.001** |
|  | 5 mg mL⁻¹ | 0.118 | 0.002 | 49.87 | **<0.001** |

Table S5: One-way ANOVA results for the growth rate parameter (r) of the different yeast species exposed to different concentrations of seaweed polysaccharides. Df: degrees of freedom. Sum Sq: sum of squares. Mean Sq: mean squares. F value: F statistic value.

| Species | Df | Sum Sq | Mean Sq | F value | p.value |
| --- | --- | --- | --- | --- | --- |
| *C. albicans* | 4 | 0,008 | 0,002 | 23,71 | **<0,001** |
|  | 10 | 0,0009 | 0,00009 |  |  |
| *C. auris* | 4 | 0,007 | 0,001 | 39,99 | **<0,001** |
|  | 10 | 0,0004 | 0,00004 |  |  |
| *C. bacillisporus* | 4 | 0,001 | <0.001 | 18,6 | **<0,001** |
|  | 10 | 0,0002 | <0,001 |  |  |
| *Cr. deuterogatti* | 4 | 0,004 | 0,001 | 51,08 | **<0,001** |
|  | 10 | 0,0002 | 0,00002 |  |  |

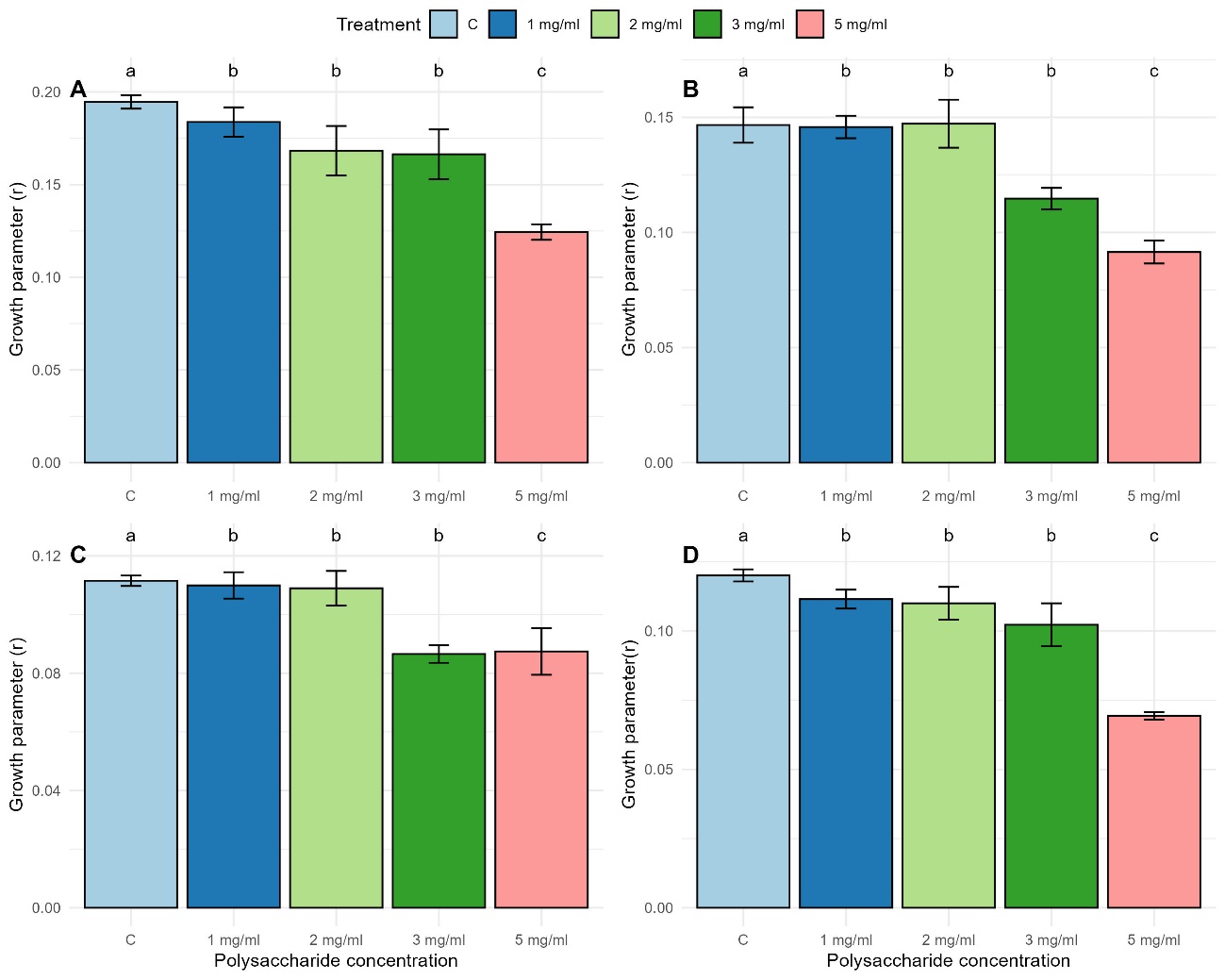

Figure S12. Effect of seaweed polysaccharides on the growth parameter (r) of A) *C. albicans*, B) *C. auris*, C) *Cr. bacillisporus*, and D) *Cr.deuterogatti* at different concentrations (1, 2, 3 and 5 mg mL⁻¹). C: control. Bars represent the mean ± standard deviation. Different letters indicate significant differences among treatments according to Tukey’s post hoc test (p < 0.05).

Table S6: One-way ANOVA results for the growth inhibition of the different yeast species exposed to different concentrations of seaweed polysaccharides. Df: degrees of freedom. Sum Sq: sum of squares. Mean Sq: mean squares. F value: F statistic value.

| Species | Df | Sum Sq | Mean Sq | F value | p.value |
| --- | --- | --- | --- | --- | --- |
| *C. albicans* | 3 | 4794 | 1598 | 72.69 | **<0.001** |
|  | 8 | 176 | 22 |  |  |
| *C. auris* | 3 | 10357 | 3452 | 106.8 | **<0.001** |
|  | 8 | 259 | 32 |  |  |
| *C. bacillisporus* | 3 | 2188.9 | 729.6 | 6.381 | **0.016** |
|  | 8 | 914.7 | 114.3 |  |  |
| *Cr. deuterogatti* | 3 | 4486 | 1495.3 | 23.52 | **<0.001** |
|  | 8 | 509 | 63.6 |  |  |

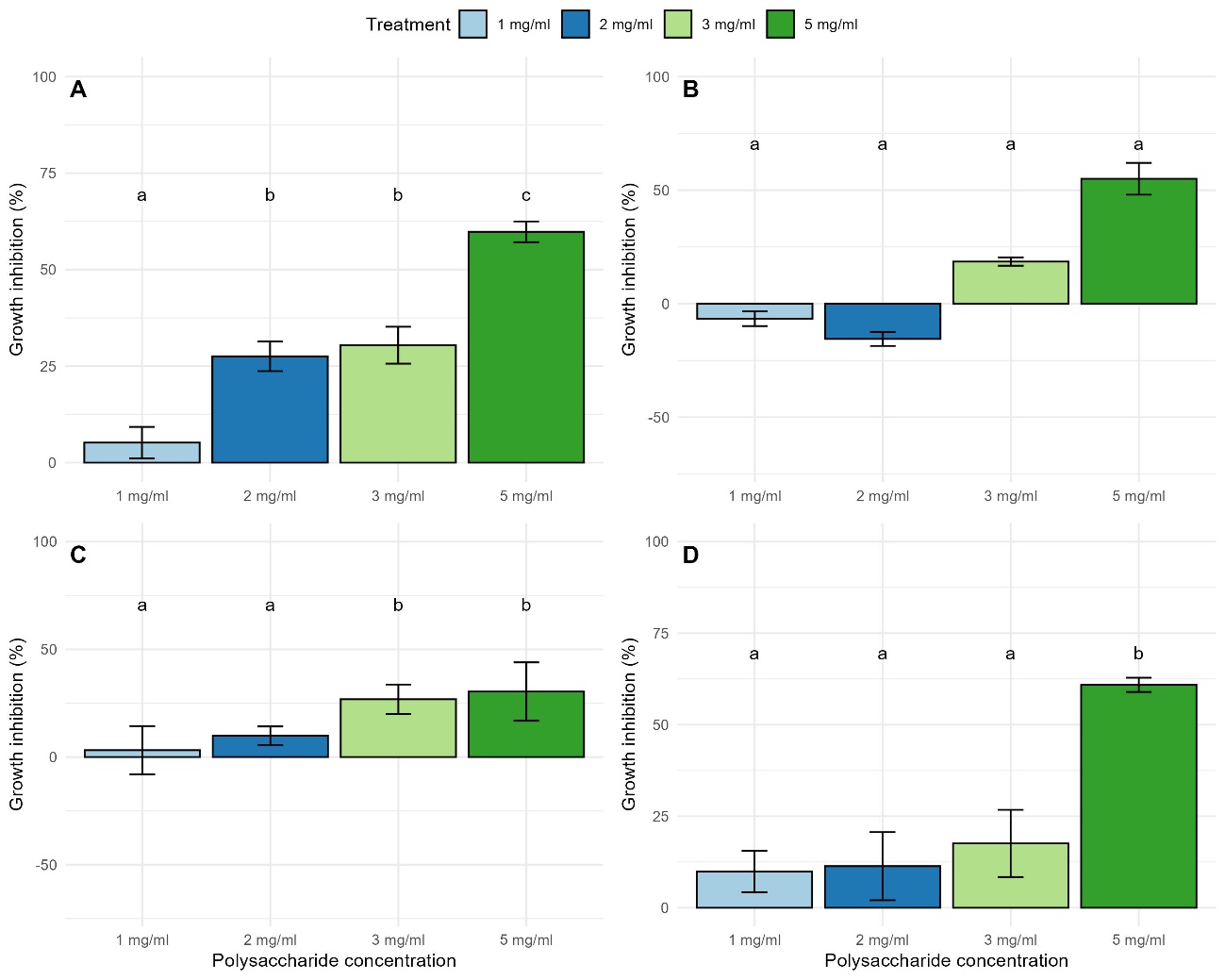

Figure S13. Effect of seaweed polysaccharides on growth inhibition (%) of A) *C. albicans*, B) *C. auris*, C) *Cr. bacillisporus* and D) *Cr. deuterogatti* at different concentrations (1, 2, 3 and 5 mg mL). Bars represent the mean ± standard deviation. Different letters indicate significant differences among treatments according to Tukey’s post hoc test (p < 0.05).
